# Functional substitutability of native herbivores by livestock for soil carbon depends on microbial decomposers

**DOI:** 10.1101/2022.02.07.479355

**Authors:** Shamik Roy, Dilip GT Naidu, Sumanta Bagchi

## Abstract

Grazing by large mammalian herbivores impacts climate as it can favor the size and stability of a large carbon (C) pool in soil. As native herbivores in the world’s grasslands, steppes, and savannas are progressively being displaced by livestock, it is important to ask whether livestock can emulate the functional roles of their native counterparts. While livestock and native herbivores can have remarkable similarity in their traits, they can differ greatly in their impacts on vegetation composition which can affect soil-C. It is uncertain how their similarities and differences impact soil-C via their influence on microbial decomposers. We test competing alternative hypotheses with a replicated, long-term, landscape-level, grazing-exclusion experiment to ask whether livestock in the Trans-Himalayan ecosystem of northern India can match decadal-scale (2005-2016) soil-C stocks under native herbivores. We evaluate multiple lines of evidence from 17 variables that influence soil-C (quantity and quality of C-input from plants, microbial biomass and metabolism, microbial community composition, veterinary antibiotics in soil), and asses their inter-relationships. Livestock and native herbivores differed in their effects on several soil microbial processes. Microbial carbon use efficiency (CUE) was 19% lower in soils under livestock. Compared to native herbivores, areas used by livestock contained 1.5 kg C m^−2^ less soil-C. Structural equation models showed that alongside effects arising from plants, livestock alter soil microbial communities which is detrimental for CUE, and ultimately also for soil-C. Supporting evidence pointed toward a link between veterinary antibiotics used on livestock, microbial communities, and soil-C. Overcoming the challenges of sequestrating antibiotics to minimize their potential impacts on climate, alongside microbial rewilding under livestock, may reconcile the conflicting demands from food-security and ecosystem services. Conservation of native herbivores and better management of livestock is crucial for soil-C stewardship to envision and achieve natural climate solutions.

## Introduction

Grazing by large mammalian herbivores is integral to the zoogeochemistry of the global carbon (C) cycle (Schmitz et al., 2014, 2018). Herbivores exert a strong influence on climate via their impacts on a large soil-C pool—over 500 Pg C in the world’s grasslands, steppes, and savannas (Derner et al., 2019; Liu et al., 2020; Naidu et al., 2022; Roy & Bagchi, 2022; Sitters et al., 2020; Witt et al., 2011). Long before the industrial era, humans started influencing global climate via their impacts on the distribution and abundance of large mammalian herbivores—prehistoric humans may have hunted many megafauna to extinction, and modern humans are progressively replacing native herbivores with domestic livestock across the world (Asner et al., 2004; Bar-On et al., 2018; Schowanek et al., 2021). Intact assemblages of native herbivores are now increasingly confined to parks and reserves; elsewhere, livestock have not only become the most abundant grazers, but they are also the most expansive land-use. Today, livestock biomass (10^11^ kg) exceeds that of all other mammals; they occur at high densities (>10^3^ individuals km^−2^) across more than 60 million km^2^ which contain over 295 Pg of soil-C (Bar-On et al., 2018; Brus et al., 2017; Gilbert et al., 2018). Further displacement of native herbivores seems inevitable due to steadily rising demands for livestock-products to accommodate evolving human-diets (Leclère et al., 2020; Thornton, 2010). Increasingly, it is important to ask whether livestock can maintain the functional roles of native herbivores they displace (Cromsigt et al., 2018; Lundgren et al., 2020; Malhi et al., 2016) because grazing can favor soil-C to decarbonize the atmosphere and offers an important natural climate solution (Naidu et al., 2022; Roy & Bagchi, 2022). If livestock can maintain the ecological functions of their native counterparts, then such substitutability can have implications for the future of biodiversity, ecosystem functions and services, food-security, and other social and environmental dimensions (Delabre et al., 2021; du Toit et al., 2012; Sayer et al., 2013; Veblen et al., 2016).

The extent to which livestock can approximate the functional influence of native herbivores on soil-C is important to inform and steer grazing management as a natural climate solution (Bossio et al., 2020; Cromsigt et al., 2018; Reinhart et al., 2022; P. Smith, 2014). On one hand, the opportunity for livestock to be functional surrogates of displaced native herbivores arises from the mutual similarity in their traits (Lundgren et al., 2020; Malhi et al., 2016; Schowanek et al., 2021). Analyses of key traits, such as body size, diet-choice, and fermentation type, hypothesize that livestock can be approximate surrogates for native herbivores, i.e., nearest neighbors in trait-space have similar ecological roles (Lundgren et al., 2020; Malhi et al., 2016; Schowanek et al., 2021). On the other hand, in many ecosystems, livestock differ from native herbivores in how they impact vegetation composition (Bagchi et al., 2012; Bagchi & Ritchie, 2010b; Price et al., 2022; Ratajczak et al., 2022; Wells et al., 2022). This alters the quantity and quality of C-input to soil from plants by changing the underlying distribution of plant traits (e.g., root:shoot ratio, C:N stoichiometry of leaf and litter, etc.) that is ultimately consequential for soil-C (Bagchi & Ritchie, 2010b; Laliberté & Tylianakis, 2012). While there is clarity on how herbivore assemblages influence vegetation communities (Augustine & McNaughton, 1998; Bagchi et al., 2012; Bakker et al., 2006), we have a limited understanding of the downstream consequences for soil-C that are ultimately controlled by microbial actions on soil organic matter (Falkowski et al., 2008; Fontaine et al., 2004; Roy & Bagchi, 2022; Sinsabaugh et al., 2013; Six et al., 2006). For instance, differences in aboveground responses may percolate belowground to alter the labile and recalcitrant fractions of soil organic matter (Bardgett & Wardle, 2003), and depending on the direction and magnitude, this can either promote or deplete soil-C (Fontaine et al., 2004; Kuzyakov, 2010). Further, livestock also release veterinary antibiotics into soil through their dung and urine which can alter soil microbial communities (Lucas et al., 2021; Wepking et al., 2019). But we do not know whether soil microbes respond differently to grazing by livestock and native herbivores, and how this influences soil-C under the two types of grazers. In other words, though the promise of functional substitutability between livestock and native herbivores might be constrained by aboveground responses by plants (Bagchi et al., 2012; Bagchi & Ritchie, 2010b), it remains tangled with belowground responses over how soil microbes respond to the two types of grazers. This limits our ability to envision and derive natural climate solutions from livestock (Bossio et al., 2020; Cromsigt et al., 2018; Reinhart et al., 2022; P. Smith, 2014) across a large fraction of the earth’s terrestrial surface (Asner et al., 2004; Bar-On et al., 2018).

Here we ask how livestock and native herbivores influence soil-C via their effects on soil microbial processes in addition to their aboveground effects (Falkowski et al., 2008; Roy & Bagchi, 2022; Sinsabaugh et al., 2016). We assess their functional substitutability in Spiti region of northern India, which is part of the greater Trans-Himalayan ecosystem spread across the Tibetan plateau and surrounding mountains (Fig. S1). We investigate long-term (decadal-scale) soil-C stocks under livestock and native herbivores, and their influence on key microbial decomposer functions related to C-cycling in soil. We take advantage of several key features in the Trans-Himalaya: (a) the livestock are a multi-species assemblage that show considerable trait-overlap with the native herbivores (Fig. 1), (b) livestock and native herbivores occupy replicate juxtaposed watersheds with comparable environmental settings (e.g., similarity in edaphic factors, climatic conditions, Fig. S1-S4), (c) their abundances are comparable such that grazing pressure on vegetation is also similar (Fig. S5). Absent major confounding influences, the livestock and native herbivore assemblages in Trans-Himalaya can help evaluate unresolved questions over how they influence soil-C (Cromsigt et al., 2018; Lundgren et al., 2020; Malhi et al., 2016; Schowanek et al., 2021).

**Figure 1.**
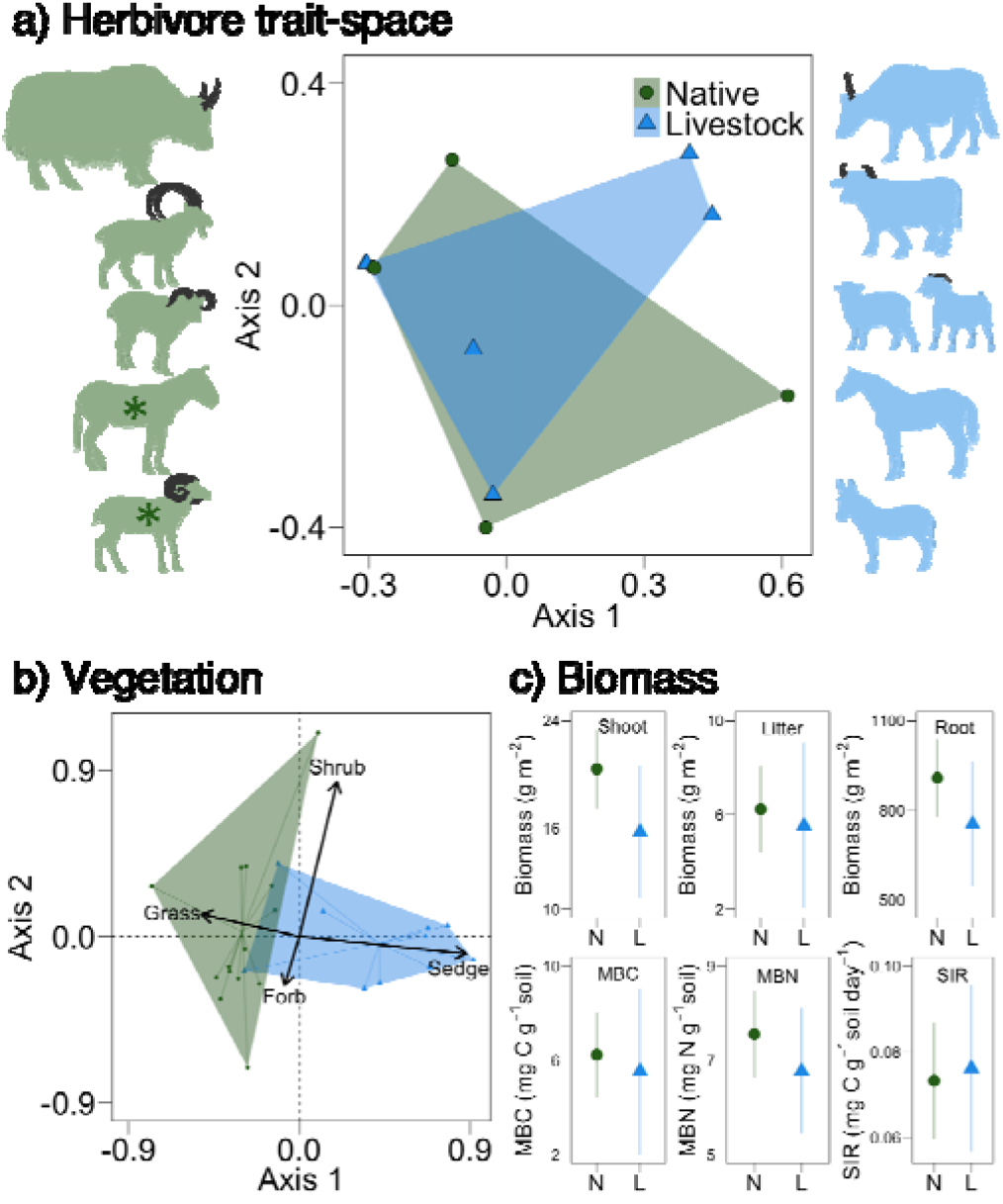
Similarities and differences between livestock and native herbivores in Spiti, India. Livestock (L: blue; yak-cattle hybrids, cattle, goat, sheep, horse, and donkey) and native herbivores (N: green, yak ibex, bharal, kiang, and argali) show considerable overlap in multidimensional trait space (a). Asterix indicates that kiang and argali are found nearby our study area in Spiti. Trait-space consists of six key traits: body mass, dietary guild, fermentation type, sexual-dimorphism, graminoid consumption, and limb morphology (Lundgren et al., 2020) with Principal Coordinates Analysis on Gower’s distance matrix (PCoA). Vegetation composition summarised with NMDS ordination on Bray-Curtis dissimilarity (b) shows that watersheds under native watersheds are dominated by grass, whereas sedges dominate under livestock. Peak plant biomass (shoot, litter and root), microbial biomass, and potential respiration (c) in native and livestock watersheds. Data are from previous studies (Bagchi et al., 2017; Bagchi & Ritchie, 2010a).

## Materials and methods

### STUDY AREA

Climate in Spiti region of Trans-Himalaya is cold and semi-arid with temperature ranging from 25 °C in summer to below −30 °C in winters (Fig. S1, S2). Precipitation occurs as winter-snow (100-200 cm yr^−1^) and summer-rain (150-300 mm yr^−1^), and vegetation growth-season is short (May-August, Fig. S2), peak biomass in July-August. Vegetation in this treeless ecosystem consists of grasses (*Poa*, *Elymus*, *Festuca*), sedges (*Carex*, *Kobresia*), forbs (*Lindelofia*, *Astragalus*) and shrubs (*Artemisia*, *Caragana*). Traditional pastoral livestock production is the major land-use across much of the Trans-Himalaya. Watersheds near village Kibber in Spiti are used by a multi-species livestock assemblage, and these are juxtaposed with watersheds that retain native herbivores (Fig. 1, Fig. S1). Natural terrain and barriers (canyons, escarpments, rivers, high ridges) restrict frequent animal movement and maintain replicate watersheds under two alternative herbivore assemblages (Fig. S1)—some used primarily by native herbivores and others by livestock (Bagchi & Ritchie, 2010b). Like other parts of the world (Veblen et al., 2016; Western et al., 2020), local extinctions of native herbivores and their replacement by livestock over the past decades remains a major conservation challenge in the Trans-Himalaya (Mishra et al., 2002; Namgail et al., 2013). The extant native herbivores are ibex (*Capra sibirica*), bharal (*Pseudois nayaur*), and yak (*Bos grunniens grunniens*) that are related to wild yak (Bos grunniens mutus; also known as B. grunniens and B. mutus, respectively; Leslie & Schaller, 2009). Currently, kiang (*Equus kiang*) are known from the fringes of the study area (Fig. S1), and there are sporadic reports of argali (*Ovis ammon*). But Tibetan antelope (*Pantholops hodgsonii*) and Tibetan gazelle (*Procapra picticaudata*) are no longer found in the study area (Mishra et al., 2002; Namgail et al., 2013). The livestock consist of yak-cattle hybrids, cattle goat, sheep, horse, and donkey (Fig. 1).

### NATIVE HERBIVORES AND LIVESTOCK OF SPITI

Cumulative biomass of native herbivores (c. 1.1×10^5^ kg) has remained comparable to that of livestock over the past few decades (c. 1.2×10^5^ kg) in the watersheds covering c. 40 km^2^ near village Kibber (Bagchi & Ritchie, 2010b; Singh et al., 2015) (Fig. S1). The livestock and native herbivores (after including kiang and argali across the broader region) show sufficiently high overlap in key traits (Lundgren et al., 2020), such as their body mass, dietary guild and graminoid consumption, fermentation type, sexual-dimorphism, and limb morphology (Fig. 1). Since the native herbivores appear to have a livestock counterpart (Fig. 1), one expects the two assemblages to be functionally similar. Yet, consistent with other parts of the world, the livestock and native herbivores differ in their impacts on vegetation composition (Bagchi et al., 2012; Bagchi & Ritchie, 2010b; Price et al., 2022; Ratajczak et al., 2022; Wells et al., 2022). Specifically, native herbivores lead to forb-and-grass dominated vegetation whereas sedges dominate under livestock (Fig. 1). Such differences in vegetation composition can be attributed to diet selectivity of the two herbivore types (Augustine & McNaughton, 1998; Bagchi et al., 2012; Bagchi & Ritchie, 2010b; Ratajczak et al., 2022). However, peak-season live biomass, both above and below-ground, are comparable (Fig. 1). In addition, standing litter biomass, and microbial biomass in soil are also comparable (Fig. 1). The two types of watersheds are also similar in several key abiotic variables (Fig. S3-S4). Soil pH is near-neutral to alkaline in both native and livestock watersheds; electrical conductivity and bulk density are also comparable (Fig. S3). Soils in both types of watersheds have sandy-loam texture (Fig. S4).

These similarities and differences can determine whether livestock can match long-term soil-C stocks under native herbivores, or not. Specifically, this not only depends on how the two types of herbivores influence C-input from plants to soil, and but also how they influence microbial processes during decomposition of soil organic matter (Bardgett & Wardle, 2003; Falkowski et al., 2008; Roy & Bagchi, 2022; Sinsabaugh et al., 2016).

### EXPERIMENTAL DESIGN

Humans arrived in Spiti in the pre-historic age (Bellezza, 2017), and livestock replaced the native herbivores from these watersheds few decades before our study began (Mishra et al., 2002; Namgail et al., 2013). Thus, *sensu stricto*, a contemporary landscape-level comparison of these watersheds cannot distinguish the effects of change in animal assemblage from any pre-existing differences between them. Therefore, instead of relying solely on prevailing differences in above- and belowground factors that influence soil-C, we determine whether any observed difference in soil-C between the watersheds can be attributed to how plants and microbes respond to grazing by the alternative herbivore assemblages. We use long-term experimental herbivore-exclusion in the replicate watersheds to quantify grazer-effects on different variables to investigate (a) whether grazing by livestock and native herbivores has different influence on soil-C, and (b) whether any difference in soil-C can be attributed to how plants and microbes respond to grazing by the two assemblages. This approach overlays a manipulative experiment over the natural experiment of two types of watersheds (Bagchi & Ritchie, 2010b). For this, starting in 2005, we set up experimental grazer-exclusion with replicated paired-and-adjacent control-and-fenced plots (10×10 m^2^, each). We used four watersheds used primarily by native herbivores, and another four by livestock (Fig. S1). We set up 3-4 paired control-and-fenced plots in each watershed (n=15 plots under native herbivores, and another n=15 plots under livestock), with a total of 30 paired control-and-fenced plots.

### SOIL CARBON AND PLANT BIOMASS

Over the next 12 years, we measured soil-C eight times at roughly inter-annual intervals in each paired plot across the different watersheds. We collected soils with a 5 cm diameter and 20 cm deep cores, and measured carbon content (TruSpec, Leco, USA) to estimate soil-C stocks (kg C m^−2^ up to 20 cm depth) under livestock and native herbivores; soil-depth rarely exceeds 20 cm in this ecosystem and in other similar mountainous landscapes (Bagchi & Ritchie, 2010b). We measured the difference in standing above- and belowground biomass at peak of growing season between paired fenced and grazed plots as an indicator of grazing pressure (Fig. S6). We collected live shoot biomass from 0.5×0.5 m^2^ quadrats in each paired plot. Subsequently we first sun-dried and then oven-dried the biomass samples to constant weight at 40 °C to obtain the dry-weights (g m^−2^). We separated live roots and other belowground structures such as rhizomes from the soil cores, and similarly obtained dry-weights (g m^−2^ up to 20 cm depth) (Fig. S6).

### SOIL MICROBIAL FUNCTIONS

In 2016, after up to 12 years of herbivore-exclusion, we estimated grazer-exclusion effects on soil microbial processes in the paired-adjacent fenced and control plots to assess whether they explain any observed differences in soil-C. These soil microbial variables represent different processes that collectively influence nutrient cycling in soil (Table S1). Residence-time of soil-C in such subhumid ecosystems is 10-30 years, whereas it can be in excess of 100 years in swamps and wetlands (Carvalhais et al., 2014; Raich & Schlesinger, 1992). Thus, much of the soil-C from the time of initial replacement of herbivore assemblages would have undergone considerable turnover. So, we expect our long-term grazing-exclusion experiment to reflect how the two types of herbivore assemblages influence soil-C at decadal timescales that is relevant to their functional influence on climate (Naidu et al., 2022). In this way, the grazing-exclusion experiment offers an opportunity to assess if differences in plant and microbial responses to grazing by different types of herbivores can explain how they influence soil-C.

In 2016, we estimated the response of key soil microbial functions to grazing: basal respiration (BR), potential respiration or substrate-induced respiration (SIR), broad soil microbial community profile as bacterial and fungal fractions (B and F), microbial biomass C and N (i.e., MBC and MBN), labile carbon (LC), labile nitrogen (LN), and recalcitrant carbon (RC) fractions of soil organic matter, microbial metabolic quotient (qCO_2_), and microbial carbon use efficiency (CUE). Since these microbial processes have seasonal dynamics (Bagchi et al., 2017; Roy & Bagchi, 2022), we measured them at five time-points during the growing season between May and September (Fig. S7). We followed standard laboratory protocols to measure these microbial variables (Bagchi et al., 2017; Robertson et al., 1999; Roy & Bagchi, 2022). Briefly, for BR we used an alkali-trap to incubate 4 g soil at 60% water holding capacity and measured respired CO_2_ (mg C g^−1^ soil day^−1^). Similarly, we measured SIR (mg C g^−1^ soil day^−1^) after adding 0.5% (w/v) of glucose to soil. We estimated broad microbial community profile as fungal and bacterial contributions to potential respiration under selective inhibition using Streptomycin (anti-bacterial) and Cycloheximide (anti-fungal) with respect to controls. We used chloroform fumigation-extraction to estimate MBC (mg C g^−1^ soil), and MBN (mg N g^−1^ soil). We measured labile and recalcitrant fractions (mg g^−1^ soil) of soil organic matter (SOM) using two-step hydrolysis with H_2_SO_4_. We estimated qCO_2_ as the ratio of BR and MBC (respiration per unit biomass, hr^−1^). We estimated CUE from established relationships between key microbial extracellular enzymes (carbon-acquiring β-glucosidase, alongside nitrogen-acquiring Leucine aminopeptidase and *N*-acetyl-β-D-glucosaminidase), MBC and MBN, and labile fractions of C and N in SOM (Sinsabaugh et al., 2013, 2016). We measured activity of the three enzymes using their specific fluorogenic substrates (4-MUB-β-D-glucopyranoside for β-glucosidase, L-Leucine-7-amido-4-methylcoumarin hydrochloride for Leucine aminopeptidase, and 4-MUB-*N*-acetyl-β-D-glucosaminide for *N*-acetyl-β-D-glucosamindase) (German et al., 2011) in soils (Fig. S8). These parameters influence microbial metabolism to determine their investment in growth (*G*) and respiration (*R*), so that 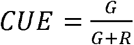 (Sinsabaugh et al., 2013, 2016). Theoretically CUE can range between 0 and 1, i.e., nil to perfect efficiency, with higher values indicating greater potential for C-sequestration. But, CUE in nature generally does not exceed 0.6 (Sinsabaugh et al., 2013, 2016).

### SOIL MICROBIAL DNA

Based on patterns seen in fungal and bacterial contribution to soil respiration, we evaluated community composition in more detail using their DNA markers from soils collected in 2019 (Fierer et al., 2005; Maestre et al., 2015). We used soils from a subset of 20 paired-adjacent plots and extracted microbial DNA (MOBIO Power Soil DNA Extraction Kit). Since fungal and bacterial fractions can have seasonal dynamics (Bagchi et al., 2017), we covered four time points during the growing season (May, July, August, and September, for a total of n = 160 samples). Next, we quantified the Internal Transcribed Spacer regions (ITS-1 and ITS-2) of rDNA for fungal abundance, and region of 16s rDNA for bacterial abundance with quantitative polymerase chain reaction, qPCR (Fierer et al., 2005; Maestre et al., 2015). We used these qPCR data to confirm whether DNA-based estimates align with respiration-based estimates of fungal and bacterial abundance. Additional details are in Appendix.

From these DNA extracts for a subset of ten paired plots, we estimated microbial taxonomic diversity in soil from Operational Taxonomic Units (OTUs). To capture seasonal community dynamics, we estimated OTUs at three time points during the growing season (May, July, and September, for a total of n = 60 samples of community wide OTUs). We amplified and sequenced the 16S rRNA hyper variable V3-V4 region in Illumina MiSeq platform, followed by taxonomic assignment using the SILVA_v138 database. Additional details are in Appendix.

### VETERINARY ANTIBIOITICS IN SOIL

Herbivore-mediated changes in plant communities, with subsequent alteration in the underlying distribution of plants traits, can influence the quantity and quality of C-input to soil (Bagchi & Ritchie, 2010a). This can lead to differences in microbial communities between livestock and native herbivores due to alteration in labile and recalcitrant fractions of soil organic matter (Bagchi & Ritchie, 2010b; Laliberté & Tylianakis, 2012). A more direct effect on microbial communities can arise from veterinary use of antibiotics on livestock that eventually enter soil via dung and urine (Albero et al., 2018; Jechalke et al., 2014; Kemper, 2008; O’Connor & Aga, 2007; Wepking et al., 2019). Livestock in Spiti are frequently treated with antibiotics such as tetracycline (therapeutic and sub-therapeutic dosage). Among the native herbivores, veterinary care for yaks is comparatively rare, while it is practically absent for ibex and bharal. So, based on the patterns seen in microbial OTUs, we quantified tetracycline (μg kg^−1^ soil) in soil from all 30 paired plots in August 2022. We used enzyme-linked immunosorbent assays (ELISA) to quantify water-extractable tetracycline levels in soil with a detection limit of 3 ppb (Immunolab, GmbH, Germany), relative to standards and controls (Aga et al., 2003; O’Connor & Aga, 2007).

### DATA ANALYSIS

We evaluated competing hypotheses (Betts et al., 2021; Tredennick et al., 2021) to determine whether the observed variation in the data merely represent spatio-temporal heterogeneity between replicates, or it is necessary to invoke grazing (fenced or grazed) and herbivore assemblage-type (livestock or native) as an explanatory variable. We used a linear mixed-effects model to evaluate long term trends in soil-C and grazer-exclusion effects on each plant and microbial variable. For soil-C we used a null-model 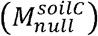 with time (years, 2006-16) and plot identity as random-effects to account for background variation due to sampling locations (plots) and times (year). This model was evaluated against the alternative competing model 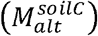 which included assemblage-type and grazing as fixed-effects, alongside the random-effects. We compared these competing models based on parsimony, goodness-of-fit, and likelihood-ratio test, i.e., AIC, RMSE, and LR (Tredennick et al., 2021).

We estimated grazer-effect on each microbial variable as the log-ratio of values in the fenced and paired-adjacent control plots i.e., 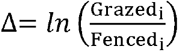 for the *i*^th^ pair, and calculated the mean and 95% CI of the log-ratio (Roy & Bagchi, 2022). As earlier, we first evaluated a null-model to assess whether the observed variation in grazer-effects can simply be attributed to spatio-temporal heterogeneity. For the null-model for the *j*^th^ microbial variable 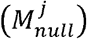, we used plot identity and month (May-September) as random-effects. In the competing alternative model for each variable 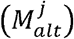, we included herbivore assemblage-type as a fixed-effect alongside the random-effects. As before, we used AIC, RMSE and LR for model comparison to judge whether herbivore assemblage-type is a necessary explanatory variable for the data (Tredennick et al., 2021). We assessed whether the data met statistical assumptions from observed and theoretical quantiles of residuals; the variables required no further transformation (Δ is already in log-scale). We used ‘nlme’ library in R (Pinheiro et al., 2020) to perform these analyses. As the data on antibiotics had many zeros (i.e., tetracycline concentration in many samples was below the detectable limit of 3 ppb), we used a generalized mixed-effects model with binomial distribution and logit-link.

We assessed the inter-relationships between grazer-effects (Δ) on different microbial variables with structural equations models (SEM). SEM help evaluate hypothesized pathways over how variation in one variable can influence another (Grace, 2006). We identified SEM paths from *a-priori* examples in the literature (Table S2). To avoid multicollinearity in the SEM arising from redundancy among the variables, we removed those variables which were derived from one another (Fig. S9). So, ΔSIR, ΔBR, ΔRC and ΔqCO_2_ were not included in this step, and we built SEM using ΔMBC, ΔMBN, ΔLN, ΔLC, ΔF, ΔB, and ΔCUE. We incorporated SEM paths as mixed-effects models for each variable as piece-wise structural equations (Lefcheck, 2016). We used directed paths (e.g., ΔF→ΔCUE) as tests of hypothesized causal relationships, and bidirectional paths (e.g., ΔMBC↔ΔMBN) for variables that are correlated or stoichiometrically coupled (Naidu et al., 2022). We did not use genetic data from ITS and 16s rDNA in the SEM because these were collected from a subset of plots, they were also not from the same year as the other microbial variables, and because estimates from e-DNA broadly matched respiration-based estimates of community composition. Similarly, we did not include tetracycline data in the SEM because when tetracycline abundance was below the detection limit of ELISA (3 ppb), we considered it to be zero—this makes Δ undefined (Δ becomes ∞). First, we evaluated inter-relationships between grazer-effects (Δ) on different microbial variables with herbivore assemblage-type as an explanatory variable. We assessed goodness of fit with Fisher’s C statistic (Roy & Bagchi, 2022; Shipley, 2009) and report standardised path coefficients alongside their statistical significance. When SEM paths are supported by data, then they offer a plausible explanation for an underlying process. When SEM paths are not supported, the hypothesized process may not give rise to the observed data.

Next, since the temporal scale and span of the microbial variables (monthly dynamics) are different from that of plant biomass and soil-C (annual or decadal dynamics), the first SEM by itself cannot directly evaluate how grazer-effects ultimately influence soil-C via microbes. While these different types of data differ in their temporal dimensions, they have a common spatial dimension (n=30 replicate pairs). So, we evaluated their connections under the expectation that effects seen on microbial processes could accumulate and potentially explain variation in soil-C. For this, we first calculated the average (mean effect on *j*^th^ variable, 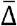) across all the months for each microbial variable, and across years for soil-C and grazing intensity, for all replicates. In this second SEM, we evaluated paths using these average values to assess whether herbivore assemblage type explains variation in soil-C via grazer-effects on microbes, alongside the effects originating from plants. As before, plot identity was a random-effect in the SEM paths as this allows the intercept to vary between samples that may have unmeasured background difference in 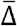 due to other factors such as variation grazing pressure, or in edaphic conditions (Naidu et al., 2022; Roy & Bagchi, 2022). We summarized and visualized the variation in microbial community composition with Principal Coordinates Analysis (PCoA) over Sorensen dissimilarity. We determined the effect of herbivore assemblage type and grazing on microbial community composition using 999 randomized iterations of the dissimilarity matrix. All analyses were performed in R 4.1.0.

## Results

### LONG-TERM TREND IN SOIL-CARBON

Model comparison showed that spatio-temporal heterogeneity was not sufficient to explain long-term variation in soil-C, and it could instead be attributed to herbivore assemblage-type. There was stronger support for 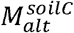 than for 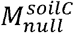 (ΔAIC=−74.39, ΔRMSE=−0.662, LR=80.386, P<0.001). However, soils under native herbivores stored 1.55±0.3 SE more carbon (kg C m^−2^) than under livestock (F_1,184_=40.11, P<0.001). Effectively, areas under livestock could realize up to 71% of the soil-C found under native herbivores (Fig. 2). Grazer-exclusion, by itself (F_1,178_=3.66, P=0.06) and as interaction (F_1,178_=1.25, P=0.26), had weak influence on soil-C (Fig. 2) even though the fenced plots received higher C-inputs from plant biomass that was not consumed by grazers (Fig. 3).

**Figure 2.**
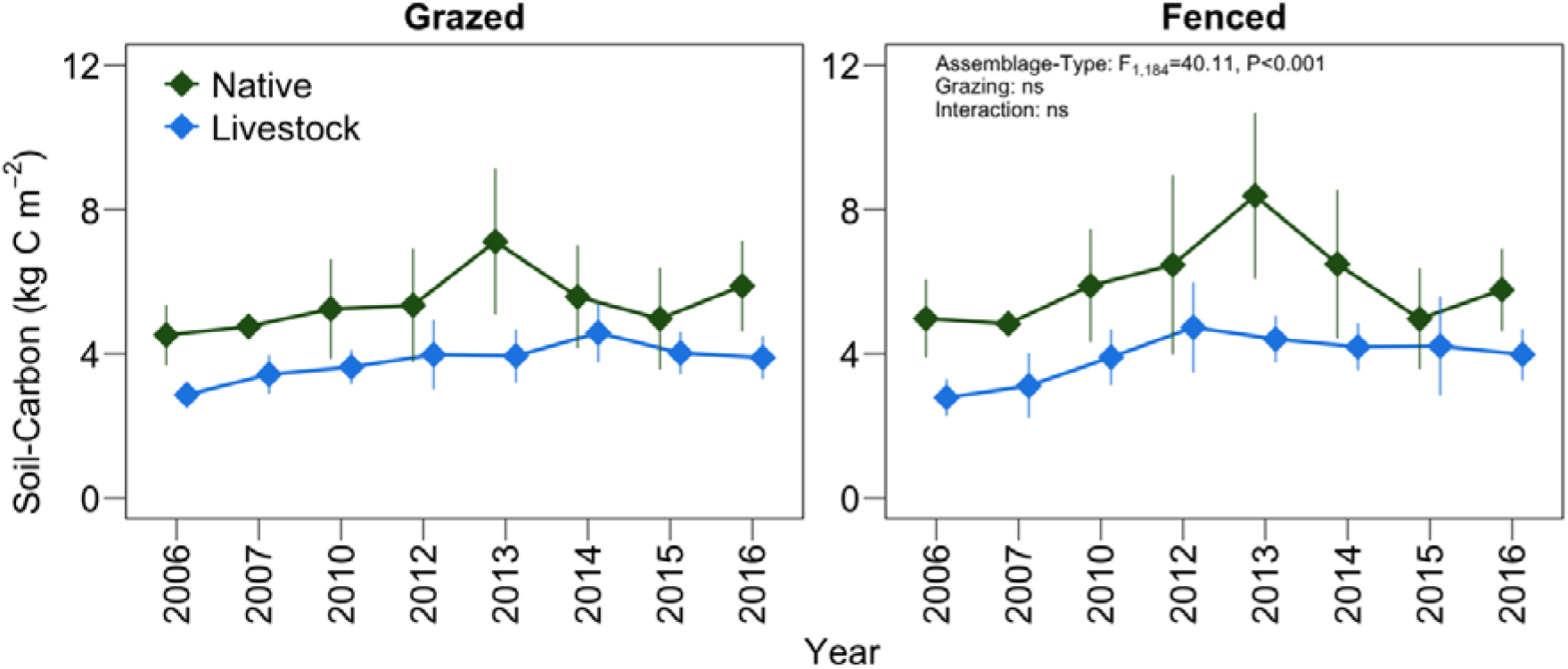
Long-term monitoring of soil-C (mean ± 95% CI) under grazed and fenced plots in watersheds with native herbivores and livestock in Spiti, India. The difference in soil-C under native herbivores and livestock was consistent for more than a decade and was not altered by grazer-exclusion. Data are from n=30 paired control (grazed) and treatment (fenced) plots.

**Figure 3.**
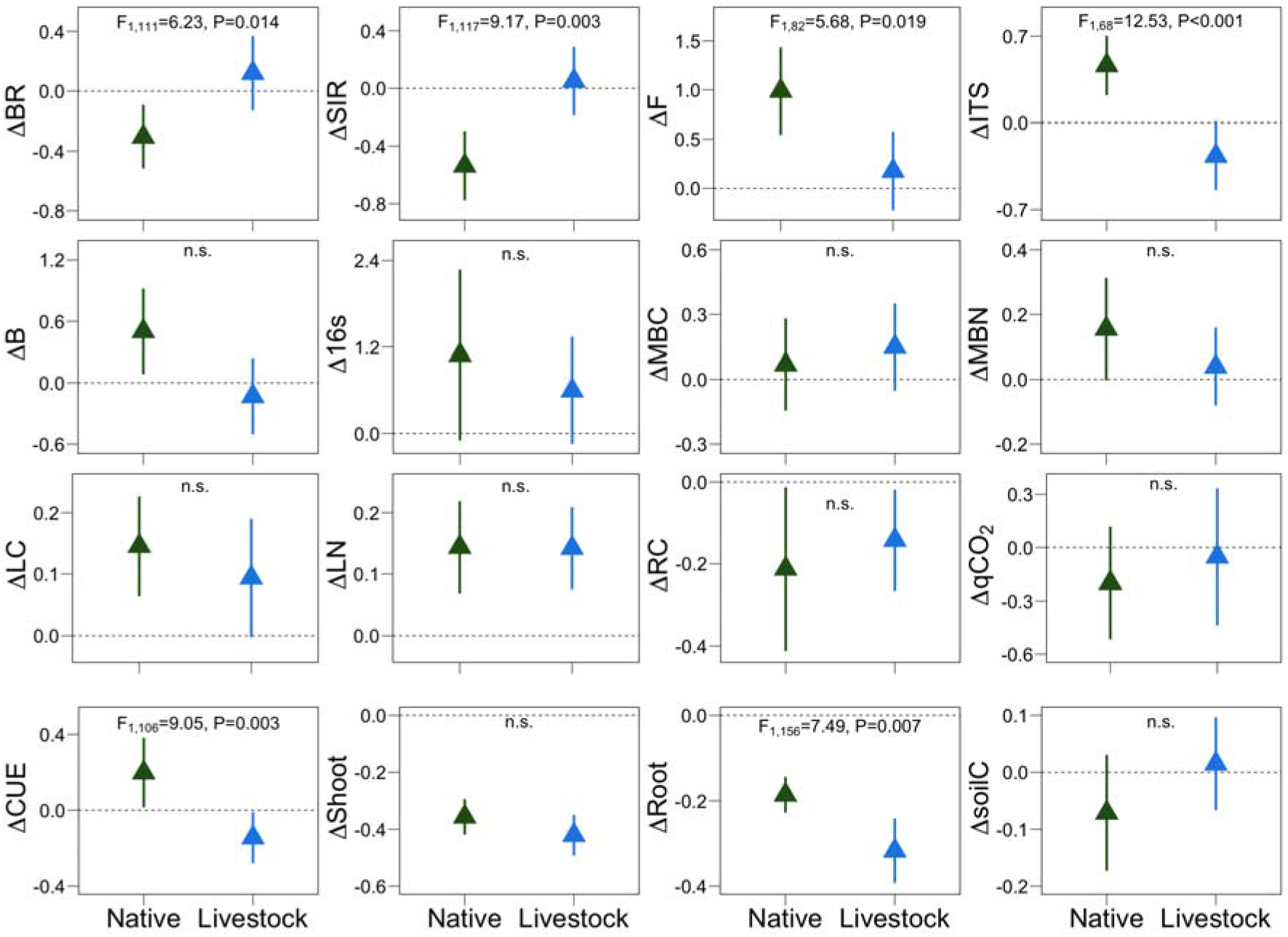
Grazer-effect (—, as mean ± 95% CI) of livestock and native herbivores on 15 plant and soil microbial variables, alongside soil-C in Spiti, India. These variables are basal microbial respiration (BR), potential microbial respiration (SIR), Fungal fraction (F), ITS DNA-amplicon abundance (ITS), Bacterial fraction (B), 16s DNA-amplicon abundance (16s), Microbial Biomass Carbon (MBC), Microbial Biomass Nitrogen (MBN), Labile-C (LC) and Labile-N (LN) fractions of soil organic matter, Microbial metabolic quotient (qCO_2_), and microbial carbon use efficiency (CUE) which were measured at monthly intervals during the growth season. Vegetation biomass (ΔShoot: from difference in peak aboveground biomass and ΔRoot: from difference in peak belowground biomass) and grazer-effect on soil-C (ΔsoilC) are from long-term decadal measurements. Direction and magnitude of differences in BR, SIR, F, ITS abundance, CUE, and Root varied with herbivore assemblage-type. But the remaining variables were not influenced by herbivore assemblage-type. Data are from n=30 permanent plots in Spiti region of northern India (except e-DNA data, which are from a subset of 20 plots). See companion Fig. S9.

### GRAZER-EFFECTS ON PLANT AND SOIL VARIABLES

Analysis of grazer-effects (grazed/fenced comparison as log-ratio Δ) indicated that the observed difference in soil-C (Fig. 2) was related to how soil microbes respond to difference in herbivore assemblage-type (whether livestock or native herbivores, Fig. 3). Livestock and native herbivores had comparable effects on B, 16s rDNA abundance, MBC, MBN, LC, LN, RC, qCO_2_, and shoot biomass (Fig. 3, Table S3). For these variables, the competing alternative models were not supported, and their variability in grazer-effects could be attributed to spatio-temporal heterogeneity rather than to herbivore assemblage-type (Fig. 3, Table S3). Experimentally removing livestock and native herbivores had comparable effects on soil-C as well (Fig. 3). But livestock and native herbivores differed in their influence on BR, SIR, F, ITS rDNA abundance, below-ground biomass, and on CUE. For these six variables, there was stronger support for the competing models than the null models (Table S3), as assemblage-type was an important explanatory variable over and beyond background spatio-temporal heterogeneity (Fig. 3). Grazing by livestock increased both basal and potential microbial respiration (BR and SIR), compared to grazing by native herbivores (Fig. 3). But grazing by livestock reduced both the fungal fraction of microbial respiration and the abundance of fungal genetic markers (F and ITS). Respiration-based estimates of fungal and bacterial fractions were consistent with DNA-based estimates (Fig. 3). Importantly, grazing by livestock reduced CUE, whereas grazing by native herbivores increased it (Fig. 3).

### INTER-RELATIONSHIPS BETWEEN SOIL MICROBIAL RESPONSES

To assess how differences in grazer-effects on one microbial variable may influence other variables, we used mixed-effects structural equation model. The model was supported by the data (SEM: Fisher’s C=4.409, *P*=0.633, df=6, AIC=66.41). This showed, that change in herbivore assemblage-type influences microbial community composition to alter CUE (Assemblage→ΔF→ΔCUE, Fig. 4). Specifically, change in herbivore assemblage from native to livestock alters the soil microbial community, which in turn is detrimental for CUE (Fig. 4). This SEM indicates that direction and magnitude of grazer-effects are consistent with higher CUE under native herbivores suggesting higher potential for net soil-C storage, compared to livestock (Fig. 2, Fig. 4).

**Figure 4.**
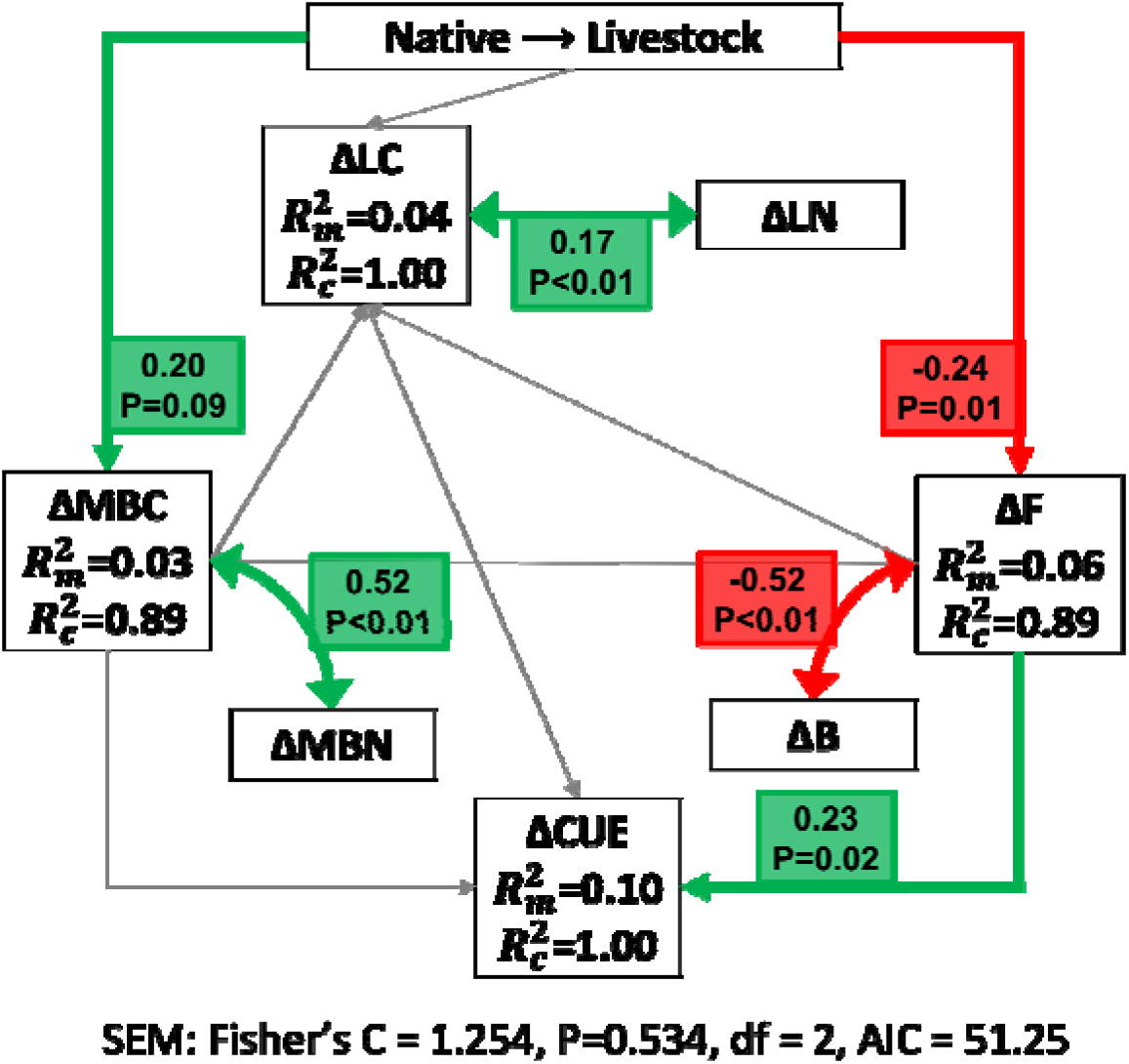
Mixed-effects structural equation model to evaluate inter-relationships between grazer-effects (—) on different microbial variables with herbivore assemblage-type as an explanatory variable. The variables are ΔMBC: microbial biomass-C, ΔMBN: microbial biomass-N, ΔLC: labile-C, ΔLN: labile-N, ΔF: fungal biomass, ΔB: bacterial biomass, ΔCUE: carbon use efficiency. SEM paths show that as herbivore-assemblage goes from native to livestock, a decrease in ΔCUE via ΔF. Unidirectional arrows represent hypothesized causal paths, bidirectional arrows indicate correlated paths, and rounded arrows indicate interactions. Standardized path coefficients and their statistical significance are shown alongside marginal and conditional R^2^ for the different variables.

### INFLUENCE OF PLANT AND MICROBIAL RESPONSES ON SOIL-C

We considered the role of herbivore assemblage-type (identity of herbivores) and vegetation biomass (shoot and root biomass), quantity and quality of C-input to soil (labile and recalcitrant pools), together, as a thought experiment to connect the inter-relationship between different microbial responses with soil-C (Fig. 5). This SEM showed that changing herbivore assemblage-type from native to livestock is detrimental to soil-C via two paths. First, herbivore assemblage-type affects soil-C through a decline in quantity of C-input to soil particularly from belowground root biomass (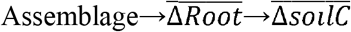, Fig. 5). Second, it also affects soil-C through effects on microbial community composition and microbial CUE (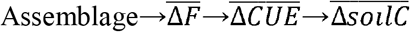; Fig. 4–5). In this way, it supported our expectation that effects arising from microbial processes at monthly scales (seen in Fig. 4) could indeed explain variation in soil-C (Fisher’s C=8.92, *P*=0.71, df=12, AIC=128.92).

**Figure 5.**
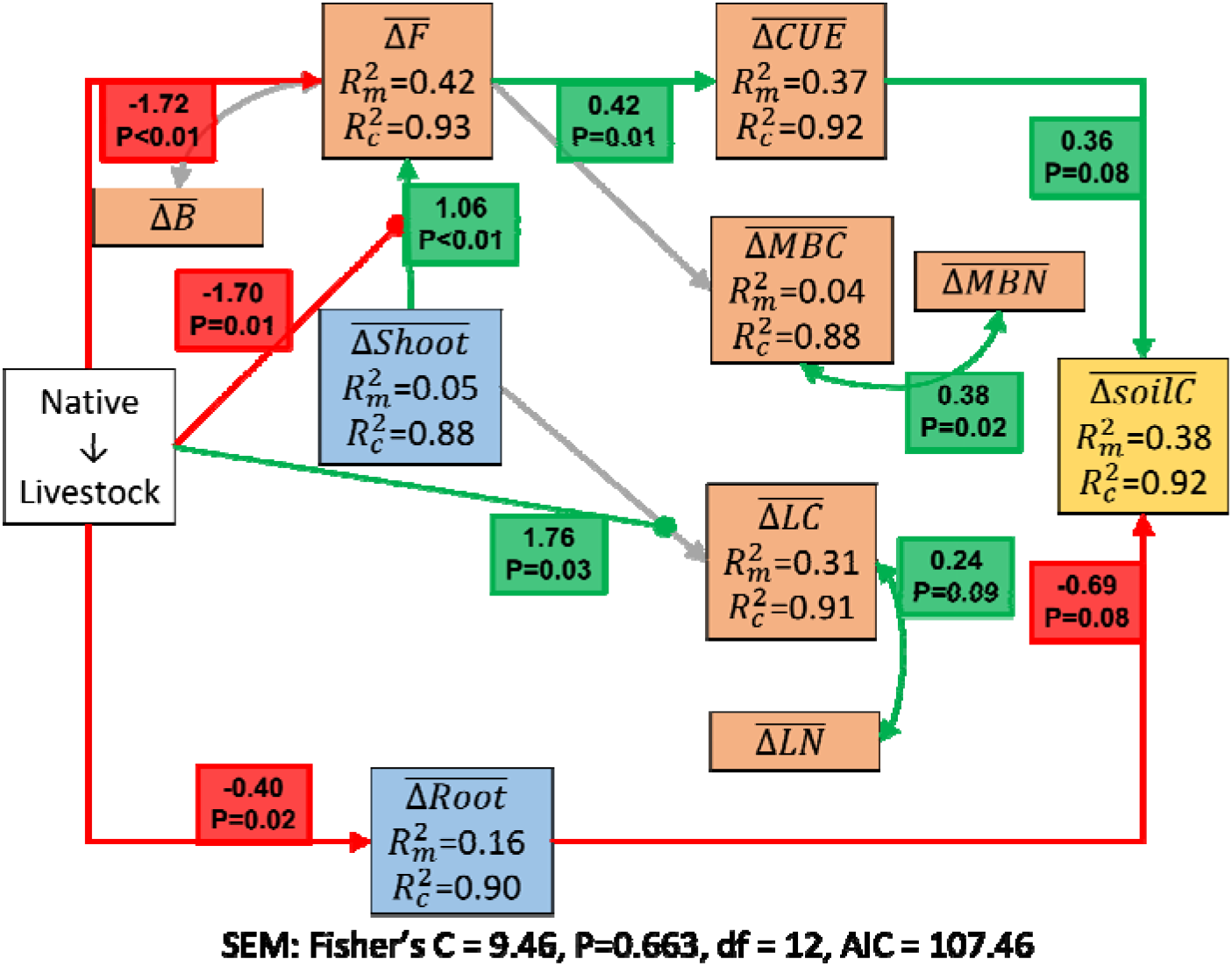
Thought experiment with mixed-effects structural equations to explore the influence of plant and microbial responses to grazing by native and livestock on soil-C. Unlike Fig. 4, here is the mean grazer-effect averaged across time (months for microbial variables, and years for soil-C and vegetation biomass). This helps evaluate the relative influence of herbivore assemblage-type, and vegetation (and), and microbial variables, on variation in soil-C. SEM paths suggest that replacing native herbivores with livestock affects soil-C in two ways: via effects originating from plants, and through a change in soil microbial communities that reduces CUE. Unidirectional arrows represent hypothesized causal paths, bidirectional arrows indicate correlated paths, and rounded arrows indicate interactions. Standardized path coefficients and their statistical significance are shown alongside marginal and conditional R^2^ for the different variables. For clarity and to reduce clutter, only a few unimportant non-significant paths are shown in grey (see companion Fig. S11 for all modelled paths).

### MICROBIAL COMMUNITIES AND VETERINARY ANTIBIOTICS IN SOIL

Following the connections between herbivores, microbes, and soil-C (Fig. 5), we evaluated microbial community composition in more detail using 16S rDNA sequencing and classification. Overall, Actinobacteriota, Acidobacteriota, Chlorofelxi, Planctomycetota, and Proteobacteria were the common and abundant taxa (Fig. S10). Microbial community composition differed between native herbivores and livestock (F_1,56_=8.03, P<0.001; Fig. 6a). However, grazer-exclusion did not alter microbial communities regardless of herbivore assemblage-type (Fig. 6a).

**Figure 6.**
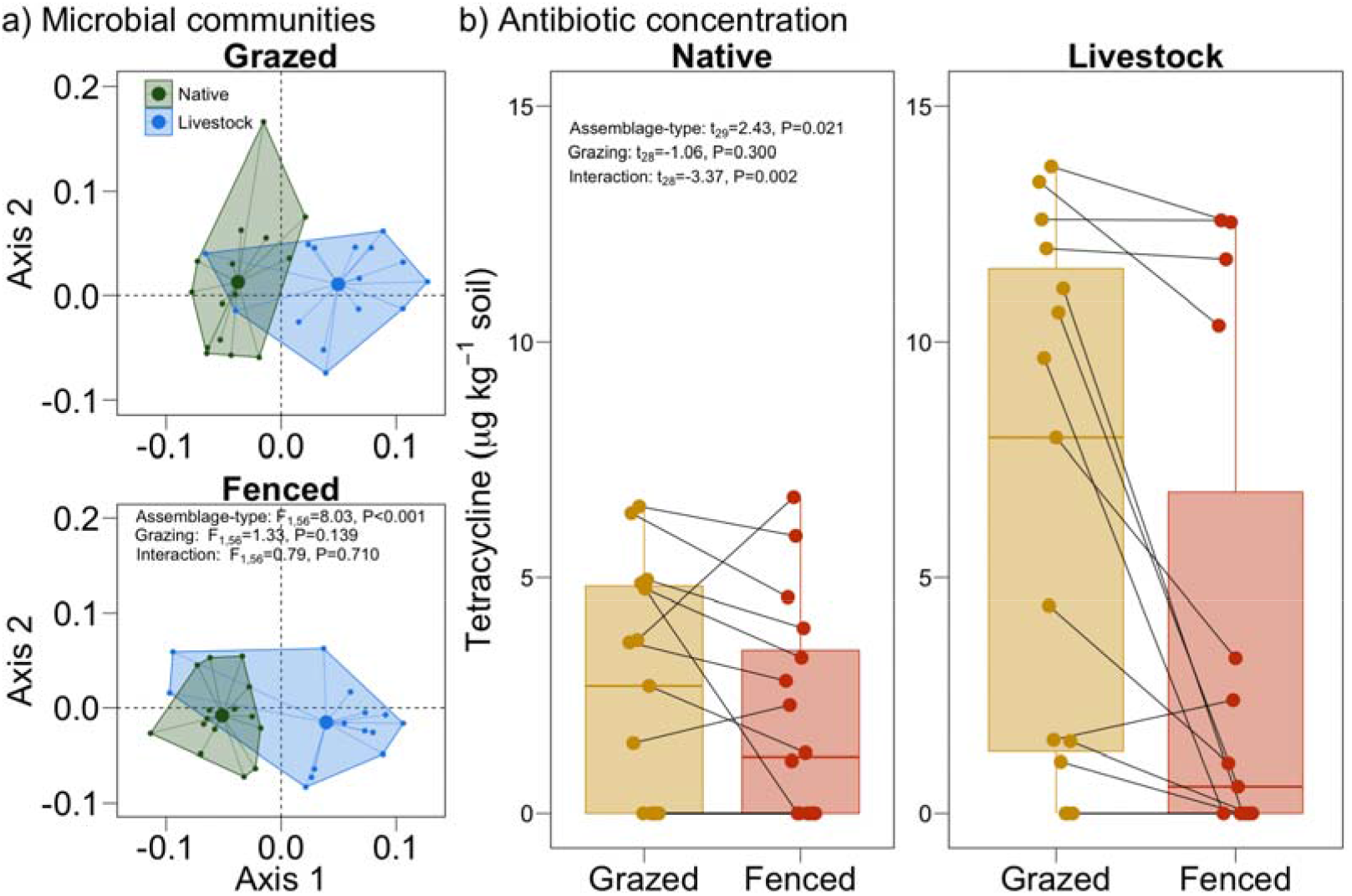
Differences in microbial community composition and antibiotic concentration across native and livestock watersheds. Difference in microbial community composition in two types of watersheds summarized with Principal Coordinates Analysis on Sorensen dissimilarity matrix of DNA-based microbial OTU data (a). Microbial communities varied between native and livestock watersheds, and there was no difference between grazed and fenced plots. Difference in antibiotic residues (tetracycline) in soil (b). Soils under livestock contain more antibiotics than soils under native herbivores, and they decline after grazer-exclusion.

To explore whether the difference in microbial communities could be related to veterinary use of antibiotics on livestock, we quantified tetracycline concentration in soil. As expected, grazed plots under livestock (mean 6.44 ± 1.42 SE, μg kg^−1^ soil) contained nearly three times more tetracycline residues than under native herbivores (2.59 ± 1.27 SE, μg kg^−1^, Fig. 6b). Expectedly, grazer-exclusion resulted in depletion of antibiotics in the fenced plots and this effect was stronger for livestock (t_28_=−3.37, P=0.002; Fig. 6b).

## Discussion

We assessed whether livestock could provide climate-mitigation services through soil-C to the same extent as the native herbivores they displace (Asner et al., 2004; Bar-On et al., 2018; du Toit et al., 2012; Schowanek et al., 2021), as this has implications for grazing management (Bossio et al., 2020; Cromsigt et al., 2018; Reinhart et al., 2022; P. Smith, 2014). We find that:- (1) multi-species herds of livestock can have remarkable functional similarities with the native assemblages they displace (Fig. 1), (2) but livestock do not emerge as perfect substitutes since they store less soil-C, compared to the native herbivores (Fig. 2), (3) differences in soil microbial responses can be the proximate explanation for why livestock are not perfect substitutes, in addition to above-ground grazer-effects that are propagated via plants (Fig. 3–5), and (4) there is supporting evidence that veterinary antibiotics can restrict soil-C under livestock (Fig. 6). These results can have implications for how we manage environmental impacts of livestock (Delabre et al., 2021; du Toit et al., 2012; Sayer et al., 2013; P. Smith, 2014; Veblen et al., 2016), and meet the rising demand for livestock-products in the coming decades (Leclère et al., 2020; Thornton, 2010).

To determine the scope of improved management of a large fraction of the earth’s terrestrial surface (Asner et al., 2004; Bar-On et al., 2018), it is important to identify the potential drivers and mechanisms that explain why livestock lag behind native herbivores in maintaining soil-C. Below we discuss explanations for the observed differences in soil-C stocks and the candidate underlying mechanisms which can be subjects for further study.

### PRE-EXISTING CONDITIONS

Since our study began after humans had already settled in the Trans-Himalayan landscape and the livestock had already displaced the native herbivores, *sensu stricto*, we cannot resolve whether livestock occupied the watersheds that were inherently low in soil-C (i.e., time_0_). Arguably, any pre-existing difference in soil-C can be maintained over subsequent decades due to comparable grazing pressures and because turnover of soil-C can occur over 10 to 30-year timescales. While pre-existing differences would explain Fig. 2, but they do not explain why livestock and native herbivores differ in their influence on soil microbial decomposers (Fig. 3). Pre-existing conditions also do not explain why the direction and magnitude of microbial responses is related to soil-C (Fig. 4–5). Alternatively, one can invoke a related argument that livestock have historically overgrazed their watersheds, thus, depleting their soil-C stocks. If this was a primary reason, then one expects soil-C to recover after cessation from grazing in the fenced plots. One also expects appreciable recovery within the decadal timescale of our study (Jones & Schmitz, 2009). However, the fenced plots in livestock watersheds did not show such a response even after 12 years of grazing-exclusion (Fig. 2). Instead, excluding native herbivores and livestock led to comparable responses in soil-C (Fig. 3). This undermines the argument that livestock watersheds are yet to recover from historical overgrazing (Fig. 2). Rather, we find that under grazing by both types of herbivores, soils effectively retain a greater fraction of their C-input from plants (Roy & Bagchi, 2022). Contrastingly in the absence of grazing, while soils receive greater C-input from plants, but are “leaky” as they fail to store soil-C, relative to the grazed plots. Such responses are known from many different ecosystems around the world and are attributable to how microbes respond to grazing (Roy & Bagchi, 2022). Finally, antibiotic residues in soil (Fig. 6) also point away from the primacy of pre-existing differences, and toward more recent factors that may have altered the microbial processes affecting soil-C. Seen together, pre-existing conditions cannot be the only explanation for the observed differences between livestock and native herbivores. Instead, it is plausible that cascading effects of antibiotics on microbial community composition and ultimately on microbial CUE play an important role, alongside the effects which originate from plants (Fig. 4–6).

### PLANT PATHWAYS

We attempt to find potential links between plant- and microbe-centric processes, above- and below-ground, and how they influence soil-C (Fig. 5). This recapitulates previous findings in terms of differences in the quantity of C-input from plants (Bagchi & Ritchie, 2010b), and accommodates downstream consequences such as quality of C-input due to changes in vegetation community composition that can alter labile and recalcitrant pools of soil organic matter (Roy & Bagchi, 2022). In other words, while the difference in vegetation composition is a determinant of the differences in soil-C, microbial responses also play an equally important role (Fig. 4–5). Unfortunately, these microbial responses do not erase the effects arising from plants and they prevent livestock from becoming perfect surrogates of the native herbivores (Fig. 4–5).

### SOIL MICROBIAL PATHWAYS

Livestock and native herbivores differ in several ways in how they influence soil microbial processes (Fig. 3). These microbial responses lead to lower CUE in soils under livestock, compared to the native herbivores (Fig. 4). CUE is a fundamental microbial trait that determines the rate at which soil organic matter is metabolized, therefore, how much C can be stored in soil (Fontaine et al., 2004; Sinsabaugh et al., 2013; Six et al., 2006). Not surprisingly, microbial community composition plays an important role because grazing by livestock is seen to reduce CUE, compared to the native herbivores (Fig. 3–4), and this can be consequential for soil-C (Fig. 5). So, the question becomes—why do livestock establish a different microbial community (Fig. 6)?

### LIVESTOCK AND SOIL MICROBIAL COMMUNITIES

Several overlapping factors can alter microbial communities under livestock, and their individual roles may not be easily disentangled from each other especially if they act in the same direction. Once established, the differences in microbial community composition may persist for long periods (López Zieher et al., 2020) due to reorganization of competition-cooperation interactions (Lechón-Alonso et al., 2021); indeed, we find them to be unresponsive to decadal-scale grazer-exclusion (Fig. 6). Differences in vegetation composition, and subsequent alteration in the quantity and quality of C-input from plants can result in microbial community differences. Additionally, veterinary care offered to the livestock releases antibiotics into soil via their dung and urine (Albero et al., 2018; Jechalke et al., 2014; Kemper, 2008; O’Connor & Aga, 2007). This can alter microbial communities towards lower CUE (Lucas et al., 2021; Wepking et al., 2019). Annually, about 10^7^ kg of antibiotics are used on livestock around the world. Within a few hours of administration, these antibiotics can enter the environment either unaltered, or as metabolites that continue to remain active (O’Connor & Aga, 2007). Indeed, soils under livestock had higher tetracycline residues (Fig. 6b). Moreover, select microbial phyla (e.g., Bacteriodota, Firmicutes, Myxococcota, Fig. S10) that are known to be particularly susceptible to antibiotics in lab-incubations (Qian et al., 2016; Shawver et al., 2021), indeed also appear to have different abundance under livestock, compared to native herbivores. This raises two unanswered questions: (1) does susceptibility to antibiotics lead to a difference in microbial community composition? And, (2) does the resultant difference in microbial community composition impact CUE to become consequential for soil-C? There is scattered evidence from laboratory cultures on how veterinary antibiotics alters soil microbial communities (Qian et al., 2016; Shawver et al., 2021), and how various microbial taxa differ in their CUE (T. P. Smith et al., 2021). However, we are yet to fully address how antibiotics can influence microbial CUE in soil in ways that ultimately also affect the global C-cycle (Lucas et al., 2021; Wepking et al., 2019). Seen together, our results identify microbial CUE as a proximate explanation for why livestock are not perfect functional substitutes for native herbivores (Fig. 3–6), to highlight the importance of soil microbial restoration in tandem with sequestering their antibiotics. In other words, livestock management may offer improved soil-C services (Bossio et al., 2020; Cromsigt et al., 2018; Reinhart et al., 2022; P. Smith, 2014) if it is coupled with microbial restoration and rewilding (Averill et al., 2022; Mills et al., 2017; Paluch et al., 2013; Wubs et al., 2016). But several questions remain unanswered on how to achieve effective microbial rewilding and whether it can sustain ecosystem-level benefits, and these can be subjected to further study.

#### MICROBIAL REWILDLING

One possible path to favor soil-C under livestock may lie in rewilding and restoring microbial communities to achieve improved ecosystem functions and services (Averill et al., 2022; Mills et al., 2017; Wubs et al., 2016). Although microbial restoration through soil-inoculations is practiced in croplands (Verbruggen et al., 2013), whether target microbial communities can establish and persist under livestock remains unknown because this problem is compounded by veterinary antibiotics. It is also unclear how we can sequester antibiotics to prevent their impact on soil-C and on global climate. Further studies on effective rewilding of microbial communities and sequestering antibiotics are needed to make livestock-production more environmentally responsible (Bossio et al., 2020; Cromsigt et al., 2018; du Toit et al., 2012; P. Smith, 2014).

In conclusion, our results highlight that multi-species and functionally diverse assemblages of livestock can partially approximate the ecological roles of native herbivores. This supports continued conservation of native herbivores and calls for new ideas to improve livestock management. Extensive therapeutic and sub-therapeutic use of antibiotics on livestock may restrict soil-C by altering soil microbial CUE. Sequestering antibiotics, alongside effective restoration and rewilding of soil microbial communities may enable livestock to emulate the services provided by native herbivores to achieve nature-based climate solutions. These may help reconcile the rising demands for livestock-products with ecosystem functions and services.

## Acknowledgements

CSIR-India provided graduate fellowship to SR; D.G.T.N. was supported by a graduate fellowship from the Divecha Centre for Climate Change; SB was supported by MHRD-India. DST-SERB, DST-FIST, DBT-IISc, ISRO-STC and MoEFCC provided funding support for fieldwork and lab-analyses. Govt. of Himachal Pradesh has supported our long-term research in Spiti. We thank Deepak Shetti, Dorje Chhering, Dorje Sheru Chhewang, Tandup Sushil Dorje, and the people of Kibber for help in the field and in the laboratory. We received generous support from Abhilash Vijay Nair, Ananya Jana, Atish Roy Chowdhury, Debarshi Dasgupta, Dipshikha Chakravortty, Ishika Pramanick, Kavita Isvaran, Praveen Karanth, Pronoy Baidya, Raju Roy, and Ritika Chatterjee. Discussions with Ben Bond-Lamberty, Mayank Kohli, and Oswald Schmitz helped us prepare the manuscript.

