## Appendix for "Functional substitutability of native herbivores by livestock for soil carbon depends on microbial decomposers"

This supplement includes:

Additional methods

Figs. S1 to S11

Table S1 to S4

Additional References

**Additional methods**

**Soil analyses**. For basal respiration (Bagchi et al., 2017; Robertson et al., 1999; Roy & Bagchi, 2022), we took 4 g of dry soil in a 200 ml plastic container. We pre-incubated the soil with 2 ml (60% water holding capacity) of water for 24 hr. Next, we added 0.5 ml of water (to eliminate the loss of water due to drying) and kept a beaker containing 1 N KOH solution in the container and then sealed. After incubation at 28 °C for 24 hr, we added 1 ml 15% BaCl2 to the KOH solution. 2 ml of KOH solution was titrated with 0.1 N HCl in the presence of phenolphthalein indicator to calculate $CO_2$ released from soil (mg C $g^{-1}$ soil $day^{-1}$). Corresponding control was water without soil. Similarly, for measuring potential microbial respiration (J. P. E. Anderson & Domsch, 1978b; Bagchi et al., 2017), protocol for basal respiration was followed except after pre-incubation instead of adding 0.5 ml water we used 0.5 ml of 2% (w/v) Glucose (net 0.5% glucose).

To measure contribution of fungi and bacteria to potential respiration we used selective inhibition of each group using anti-microbial agents (J. P. E. Anderson & Domsch, 1973; Bagchi et al., 2017). Streptomycin (anti-bacterial) and Cycloheximide (anti-fungal) was used. Both the anti-microbial agents were used at 8 mg/g soil at the beginning in the form of solution before pre-incubation, followed by the same protocol as above.

Microbial biomass was measured by chloroform fumigation and extraction (Bagchi et al., 2017; Beck et al., 1997; Jenkinson & Powlson, 1976; Robertson et al., 1999) in which 4 g soil was pre-incubated with water at 60% water holding capacity for 24 hr. Then the soils were kept in desiccator along with ethanol-free chloroform for fumigation. After fumigation, soils were aerated overnight to remove residual chloroform. 0.05 M $K_2SO_4$ was used to extract C and N from the soil, and was measured using TOC/TN analyzer (Shimadzu LCPH/CPN, Japan). Microbial biomass carbon (MBC) and microbial biomass nitrogen (MBN) was calculated by taking the difference in C and N in fumigated samples and corresponding unfumigated samples acting as control. Extraction efficiency was considered to be 0.45 for MBC and 0.54 for MBN.

We followed two-step hydrolysis with $H_2SO_4$ (Khalili et al., 2016; Rovira & Vallejo, 2002; Roy & Bagchi, 2022), to measure labile pool of SOM. Labile-C pool can be divided into two parts based on the degree of acid hydrolysis with pool-I being easily hydrolyzed and pool-II takes longer time and higher acidity to get hydrolyzed. Pool-I majorly consists of hemi-cellulose and non-cellulosic polysaccharide derived either from plant or microbes, whereas pool-II is primarily cellulose. Labile-C was taken as the sum of pool-I and pool-II. The difference between total organic carbon and labile-C was the recalcitrant-C pool.

Enzyme activity was measured following standard protocols (German et al., 2011; Roy & Bagchi, 2022; Saiya-Cork et al., 2002), where 1 g of soil was homogenized in 100 ml of Tris buffer of pH 7.8, as this is the average pH of the sampled soil to prepare a slurry. Enzyme specific substrates, namely 4-MUB beta-D-glucopyranoside for Beta-glucosidase, L-Leucine-7-amido-4-methylcoumarin hydrochloride for Leucine aminopeptidase, and 4-MUB N-acetyl-beta-D- glucosaminide for N-acetylglucosamindase, were prepared in Tris buffer of pH 7.8 at 200 mM concentration. For the assay, 200 µL of substrate was added to 50 µL of soil slurry on a microplate and incubated at 20 °C (average temperature during the growth season), for 2 hr. Reactions were stopped by adding 10 µL of 2M NaOH after incubation. Fluorogenic measurements were done in in 96-well fluorimeter (Tecan Infinite® M200 Pro, Mannedorf, Switzerland) at excitation wavelength 365 nm and emission wavelength 450 nm. Enzyme activities were calculated against their respective fluorescence standards and expressed as nmol $hr^{-1}$ $g^{-1}$ soil. For each sample, all the enzymes were assayed four times with corresponding controls, and then averaged (median) for further analysis.

We quantified ITS region for fungal abundance and 16s rRNA for bacterial abundance using qPCR in a CFX384 Touch Real-Time PCR Detection System (Bio-Rad

Laboratories Inc., Hercules, CA, USA). The primers ITS1 (5′-TCCGTAGGTGAACCTGCGG -3′) and ITS2 (5′- GCTGCGTTCTTCATCGATGC -3′) were used to estimate fungus (Fierer et al., 2005; Maestre et al., 2015). Similarly, 341f (5′-CCTACGGGAGGCAGCAG -3′) and 534r (5′- ATTACCGCGGCTGCTGGCA -3′) primers were used to estimate bacteria (Bru et al., 2008). The qPCR reaction mixture contained 4 µL of 2× KAPA SYBR® FAST qPCR master mix, 0.5 µL of each primer (10 µM), 0.5 µL of DNA template. For fungus, amplification was initiated by denaturation at 95 °C for 5 min followed by 45 cycles of 95 °C for 30 s, 56 °C for 30 s and 72 °C for 45 s. For bacteria, amplification was initiated by denaturation at 95 °C for 5 min followed by 45 cycles of 95 °C for 30 s, 68 °C for 30 s and 72 °C for 45 s. To generate a standard curve, we made serial dilutions ($10^{-1}$ to $10^{-5}$) of the genomic DNA of haploid *Saccharomyces cerevisiae* (fungi) and *Escherichia coli* (bacteria) (Shahsavari et al., 2016) and abundance were expressed as copies $g^{-1}$ soil.

Microbial OTUs were estimated following amplification and sequencing of the 16S rRNA hyper variable region V3-V4 for all the DNA samples. Briefly, 25 ng DNA was used along with KAPA HiFi HotStart Ready Mix and 100 nm final concentration of modified 341F (5’-CCTACGGGNGGCWGCAG-3’) and 785R (5’-GACTACHVGGGTATCTAATCC-3’) primers for amplification (Klindworth et al., 2013). Amplification was accomplished by initial denaturation at 95 °C for 5 min followed by 25 cycles of 95 °C for 30 s, 55 °C for 45 s and 72 °C for 30 s with a final extension at 72 °C for 7 min. Sequencing was done using Illumina MiSeq. High-quality amplicon sequences were analyzed using Mothur (Schloss et al., 2009). Briefly, for every sample, amplicon pairs were aligned with each other to form contigs, followed by quality filtering. Only contigs with length between 300 bp and 532 bp were retained, and rest were removed. To correct for the non-specific amplification of regions other than 16s bacterial rRNA gene, contigs were aligned to a known database for 16s rDNA.

Any ambiguous contigs aligning to other regions on the database were discarded. Chimeras were removed using reference-based chimera checking following UCHIME algorithm (Edgar et al., 2011). Taxonomy was assigned using SILVA_v138 database. The contigs were then clustered into OTUs (Operational Taxonomic Unit). As of 2020, $9.4 \times 10^6$ sequences are registered in the SILVA_v138 database for 16s rDNA. The most common phyla were Actinobacteriota, Acidobacteriota, Planctomycetota, Proteobacteria and Verrucomicrobiota. After the classification, OTU abundance was estimated. As an additional level of quality filtering OTUs with total abundance less than 20 in the full data set were removed. To account for the errors in the extraction process we also extracted from a negative control (no soil control). However, there was no significant amplification following PCR, and hence was not sequenced further.

Fig. S1

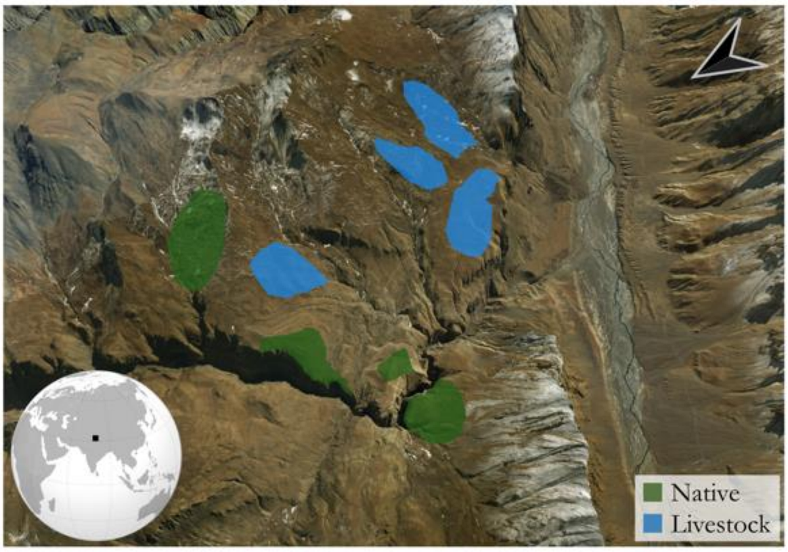


Fig. S1. Map of the study area and watersheds near village Kibber in the high-altitude region of Spiti, Trans-Himalaya, northern India. The coloured polygons represent eight watersheds spread over c. 40 $km^2$. The native and livestock watersheds are demarcated by high ridges, escarpments and ravines, that establish replicates of two types of herbivore-assemblages (dominated by either livestock or by native herbivores).

Fig. S2

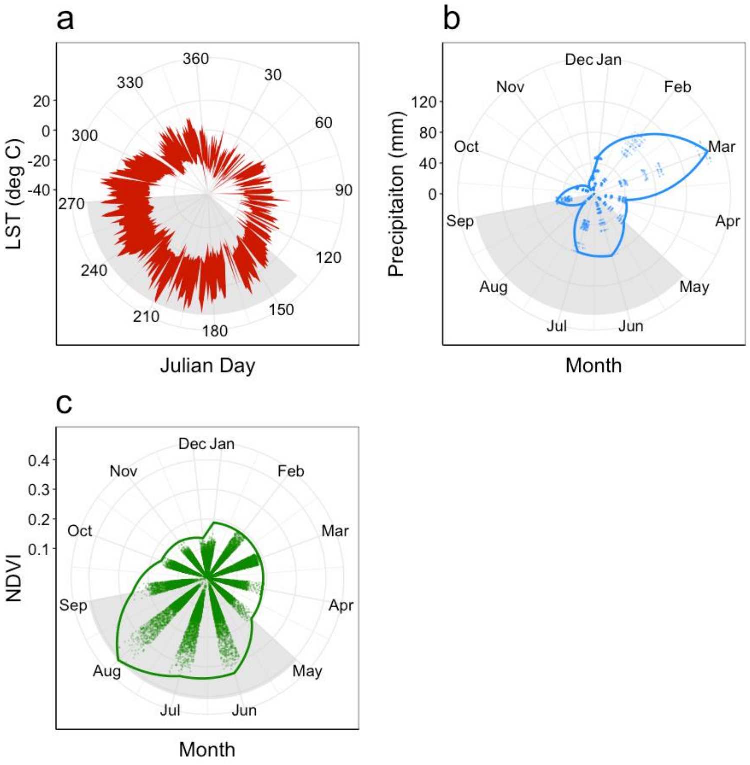


Fig. S2. Seasonality in Spiti, India, in terms of daily temperature range (a), precipitation (b), and satellite-derived normalised difference vegetation index or NDVI (c) over the last five years (2013-2017). Favourable period for vegetation growth, spanning May-August, is highlighted in grey. Temperature data at 1-km resolution are from MODIS-land surface temperature (LST), precipitation data at 5.5-km resolution are from CHIRPS, and NDVI data at 30-m resolution are from LANDSAT-7.

Fig. S3

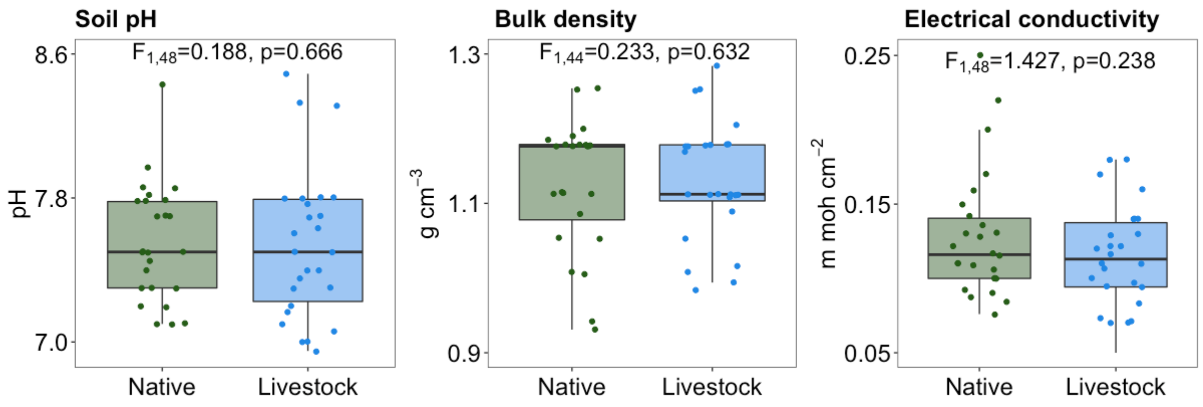


Fig. S3. Different soil edaphic factors in two types of watersheds – dominated by native or livestock, in Spiti, India. Watersheds are comparable in their edaphic conditions. Data are from previous studies (Bagchi & Ritchie, 2010b; Roy & Bagchi, 2022).

Fig. S4

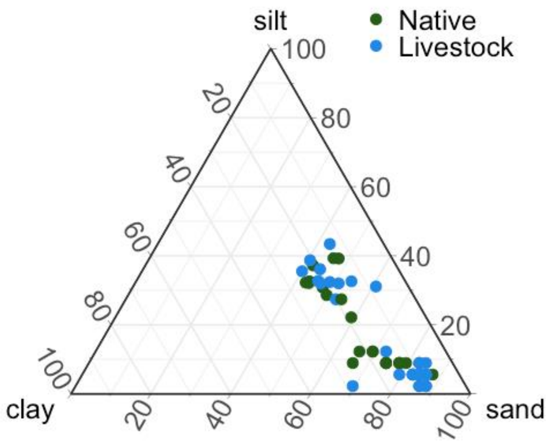


Fig. S4. Soil texture in two types of watersheds – dominated by native or livestock, in Spiti, India. Both watershed-types have sandy-loam soil texture. Data are from previous studies (Bagchi & Ritchie, 2010b; Roy & Bagchi, 2022)

Fig. S5

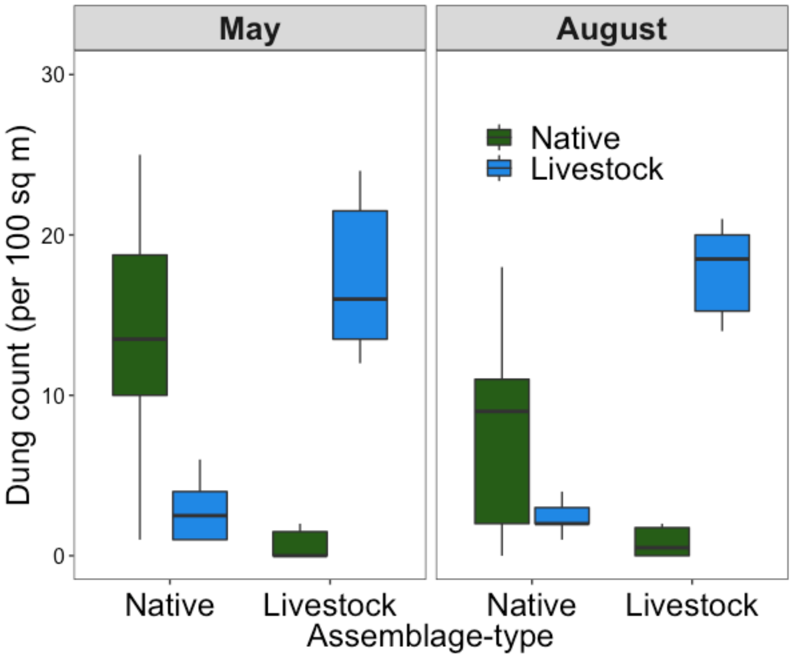


Fig. S5. Dung counts of different herbivores in two types of watersheds – dominated by wild-herbivores or livestock, in Spiti, India, during early-season (May) and late-season (August). The native herbivores are bharal, ibex, and yak. Livestock are goat, sheep, yak-cattle hybrids, horse, and donkey. Patterns suggest that native and livestock occupy separate watersheds with little habitat overlap (GLM, $z=15.48$, $p<0.001$, based on Poisson distribution). Data are from previous studies (Bagchi & Ritchie, 2010b).

Fig. S6

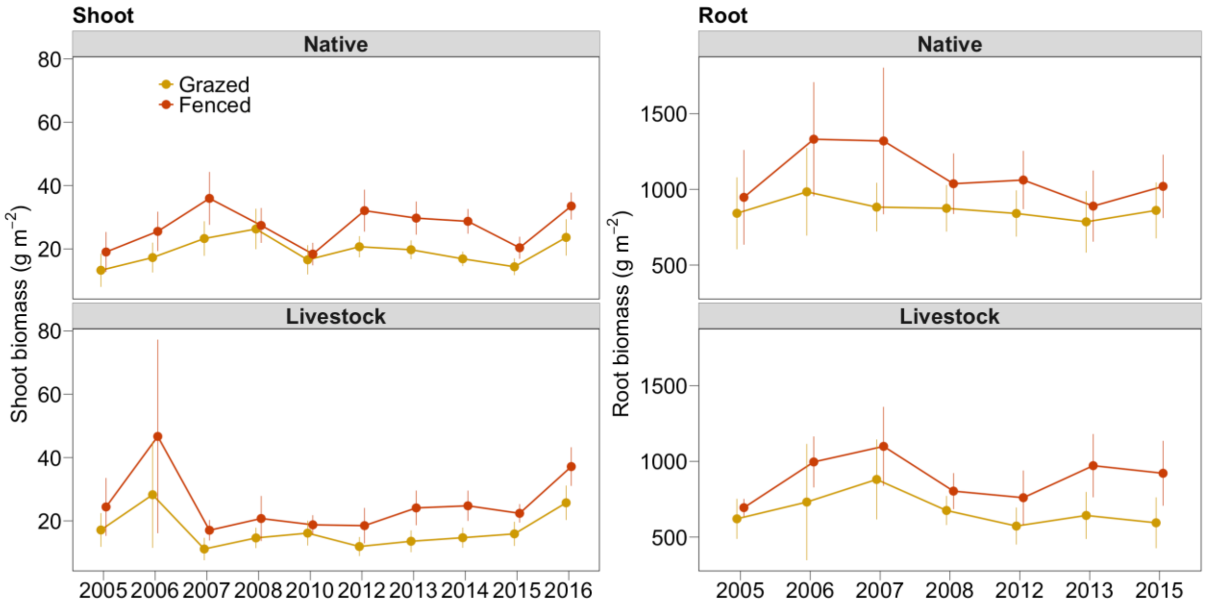


Fig. S6. Temporal variation in above- and below-ground biomass to different herbivore assemblage type in Spiti, India. Data are from n=30 paired plots (grazed and fenced) of which n= 15 paired plots are dominated by native-herbivores and n=15 paired plots are dominated by livestock.

Fig. S7

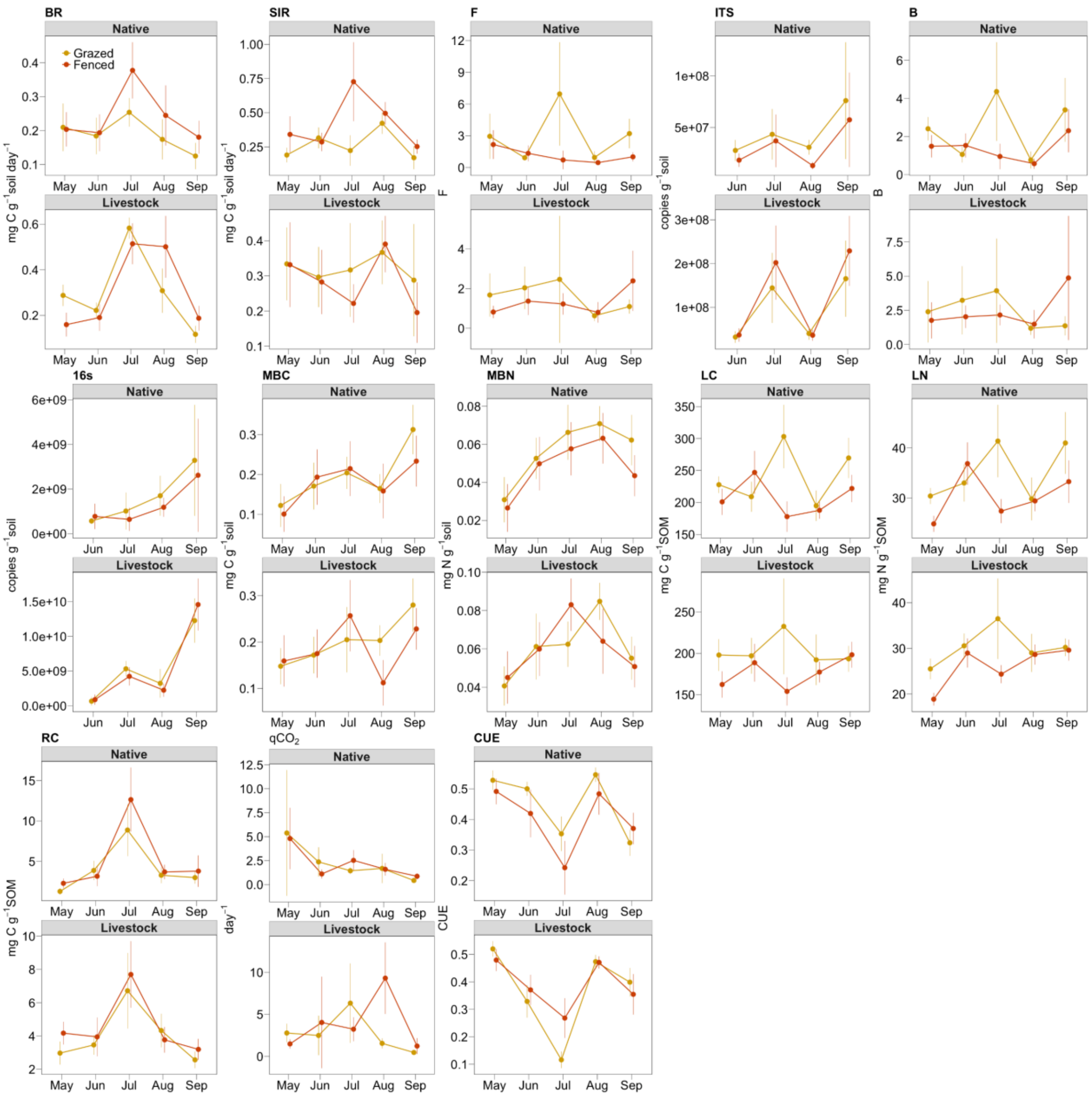


Fig. S7. Temporal variation in soil microbial responses to different herbivore assemblage type in Spiti, India. Data are from n=30 paired plots (grazed and fenced) of which n= 15 paired plots are dominated by native-herbivores and n=15 paired plots are dominated by livestock (except e-DNA data, which are from a subset of 20 plots).

Fig. S8

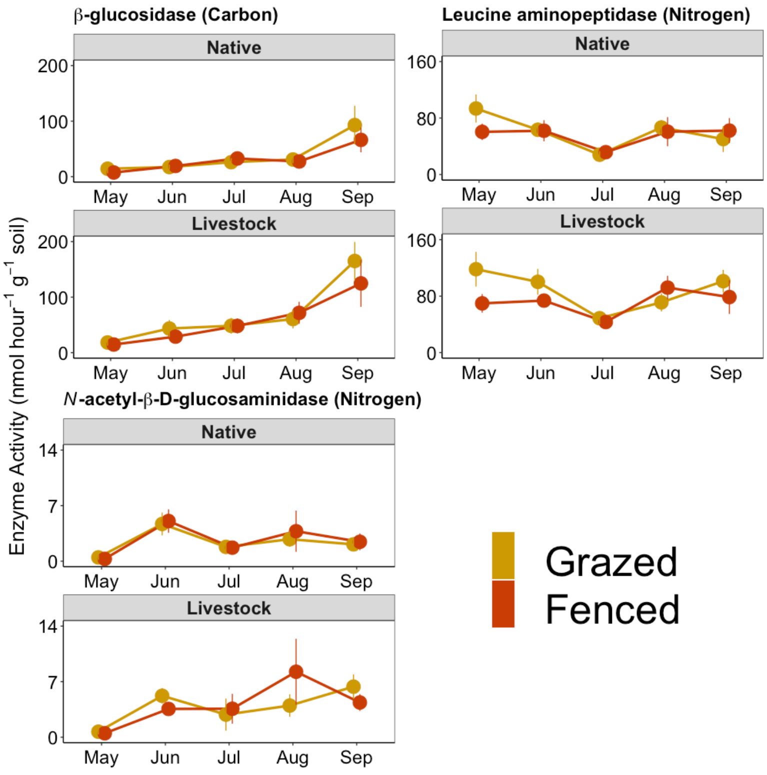


Fig. S8. Temporal variation in three soil microbial extracellular enzymes in Spiti, India. Data Sare from n=30 paired plots (grazed and fenced) of which n= 15 paired plots are dominated by native-herbivores and n=15 paired plots are dominated by livestock.

Fig. S9

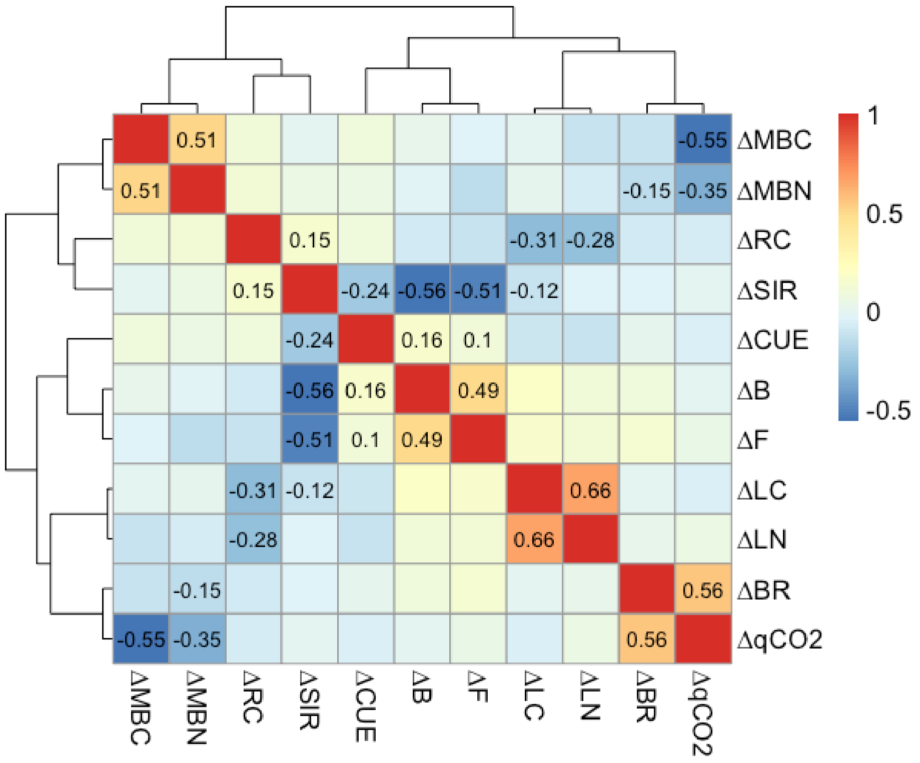


Fig. S9. Correlation between grazer-effects on interrelated soil and microbial variables. Grazer-effect (Δ, ln response-ratio) was calculated from grazed and fenced paired plots. ΔBR: basal respiration, ΔSIR: potential respiration, ΔMBC: microbial biomass-C, ΔMBN: microbial biomass-N, ΔLC: labile-C, ΔLN: labile-N, ΔRC: recalcitrant-C, $\Delta qCO_2$: Metabolic quotient, ΔF: fungal fraction, ΔB: bacterial fraction, ΔCUE: carbon use efficiency.

Fig. S10

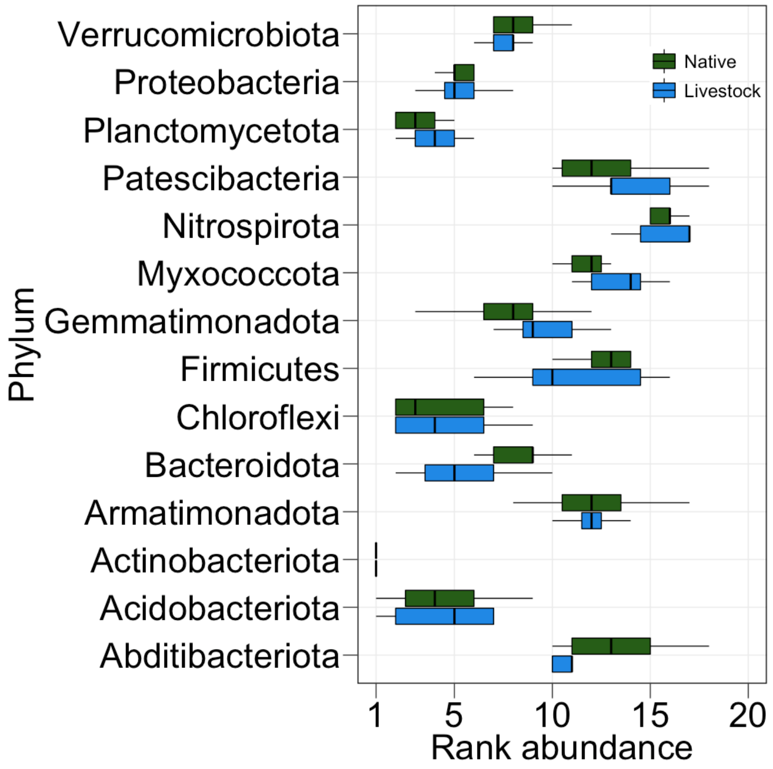


Fig. S10. Rank abundance of 14 common microbial phyla watersheds used by either native herbivores or livestock in Spiti, India. Some phyla (e.g., Bacteriodota, Firmicutes, Myxococcota) show differences, while others (e.g., Actinobacteriota, Proteobacteria) do not differ. Data are from grazed plots (5 plots × 3 months × 2 assemblage-types)

Fig. S11

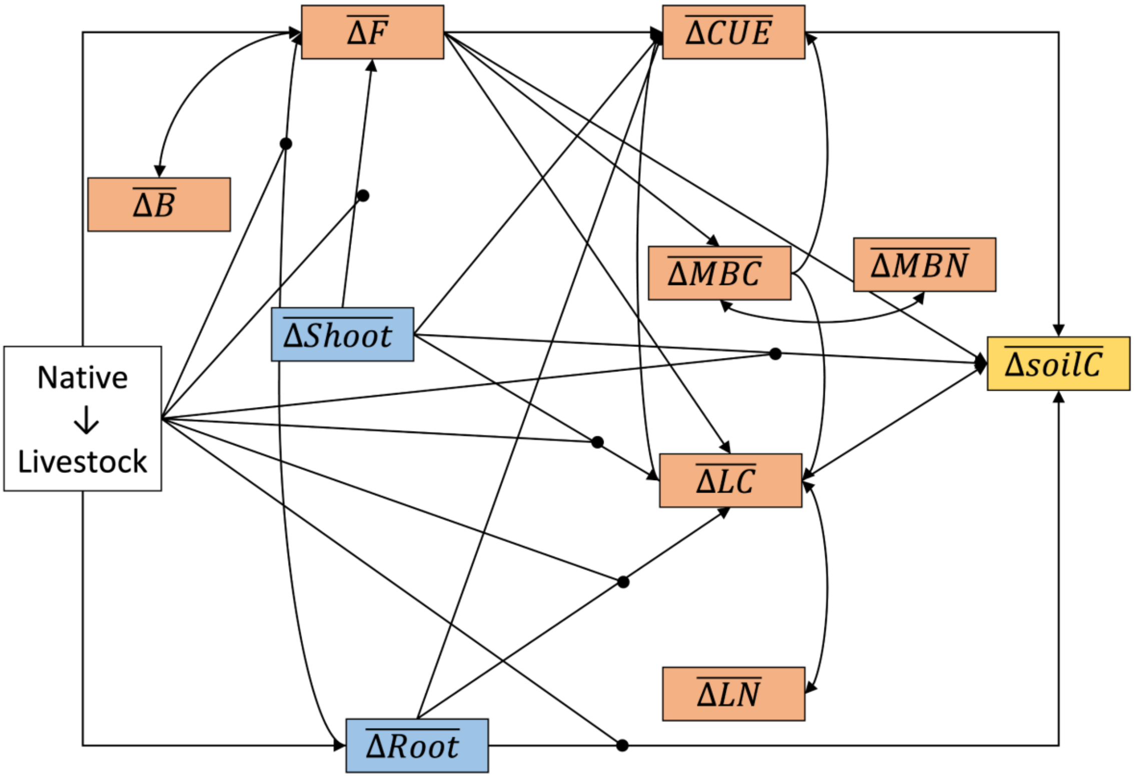


Figure S11. Companion figure for Fig. 5 with SEM showing all modeled paths. A priori examples for the hypothesized paths are summarized in Table S2.

Table S1: Summary of different soil variables

| **Soil microbial variable** | **Role in nutrient cycles** | **Reference** |
|---|---|---|
| Basal respiration | It is the baseline flux of C through soil respiration. It is an indicator of active microbial biomass and metabolism. | (J. P. E. Anderson, 1983) |
| Substrate induced respiration | It is the measure of potential microbial respiration when microbes receive easily metabolizable substrates. | (J. P. E. Anderson & Domsch, 1978a) |
| Fungal and bacterial fractions | They are measured as the contribution of active fungal and bacterial biomass towards respiration. High (or low) values for fungal and bacterial fractions represent slow (or fast) C-cycling. It is also an approximation of community composition. | (J. P. E. Anderson & Domsch, 1973, 1975) |
| Microbial biomass-C and -N | These are measures of microbial biomass in terms of mg biomass per gram soil. These represent microbial abundance as well as stoichiometry of C and N. High values of MBC and MBN indicate microbial immobilization of C and N. | (Jenkinson & Powlson, 1976) |
| Labile-C and -N | These represent the fraction of soil organic matter which can be readily metabolized by microbes and have short residence time in the soil. | (Berendse et al., 1987; Schmidt et al., 2011) |
| Recalcitrant-C | This represents the fraction of soil organic matter which cannot be readily metabolized by microbes and has long residence time in the soil. Higher recalcitrance favors long-term C-storage. | (Berendse et al., 1987; Schmidt et al., 2011) |
| qCO2 (Metabolic quotient) | It is the rate of microbial respiration per unit microbial biomass. It influences the rates of C cycling in soil. Higher qCO2 corresponds to greater C allocation for growth than to respiration, thereby stabilizing soil-C. | (T. Anderson & Domsch, 1993; Pirt, 1975) |
| Carbon use efficiency (CUE) | It is the ratio of C allocated to growth and assimilation. This is a fundamental microbial trait which determines the rate of C-cycling and storage in soil. | (Sterner & Elser, 2002) |

Table S2: A-priori evidence in literature for modelling hypothesized paths in SEM.

| **Path** | **References** |
|---|---|
| Herbivore assemblage-type → MBC | (Wang et al., 2018) |
| Herbivore assemblage-type → LC | (Wang et al., 2018) |
| Herbivore assemblage-type → F | (Bardgett & Wardle, 2003) |
| MBC → LC | (Xu et al., 2021) |
| MBC → CUE | (Cotrufo et al., 2013; Sinsabaugh et al., 2013) |
| MBC → MBN | (Heuck et al., 2015) |
| MBN → MBC | (Heuck et al., 2015) |
| F → MBC | (de Vries et al., 2006; Waring et al., 2013) |
| F → CUE | (Six et al., 2006) |
| F → LC | (Carney et al., 2007; Malik et al., 2016) |
| F → B | (de Boer et al., 2005) |
| B → F | (de Boer et al., 2005) |
| LC → CUE | (Sinsabaugh et al., 2013) |
| LC → LN | (Naidu et al., 2022) |
| LN → LC | (Naidu et al., 2022) |
| Herbivore assemblage-type → Shoot | (Bagchi & Ritchie, 2010a, 2010b) |
| Herbivore assemblage-type → Root | (Bagchi & Ritchie, 2010a, 2010b) |
| Herbivore assemblage-type → soilC | (Bagchi & Ritchie, 2010b) |
| Root → F | (Eisenhauer et al., 2017) |
| Root → soilC | (Bagchi & Ritchie, 2010b; McSherry & Ritchie, 2013; Schmidt et al., 2011) |
| Shoot → F | (Navrátilová et al., 2019) |
| Shoot → soilC | (Bagchi & Ritchie, 2010b; McSherry & Ritchie, 2013) |
| CUE → soilC | (Fontaine et al., 2004) |

| | |
|---|---|
| Root → LC | (Bardgett & Wardle, 2003; Peixoto et al., 2021; Rossi et al., 2020) |
| Root → CUE | (Cotrufo et al., 2013) |
| Shoot → CUE | (Cotrufo et al., 2013) |
| Shoot → LC | (Bardgett & Wardle, 2003; Jackson et al., 2017; Lajtha et al., 2014) |
| F → soilC | (Jastrow et al., 2007; Kallenbach et al., 2015; Strickland & Rousk, 2010) |
| LC → soilC | (Naidu et al., 2022; Schmidt et al., 2011) |

Table: S3. Model comparison of two linear models $M_0$ (null model) and $M_A$ (alternating competitive model) for soil-C, 13 microbial variables, and vegetation biomass. Grazer-effect ($\Delta = ln\left(\frac{grazed}{fenced}\right)$) was calculated from grazed and fenced paired plots

| | Null model ($M_{null}$) | | Competing model ($M_{alt}$) | | Model comparison | | | |
|---|---|---|---|---|---|---|---|---|
| | AIC | RMSE | AIC | RMSE | ΔAIC | ΔRMSE | Likelihood ratio | P-value |
| **Soil-C** | **6855.44** | **2230.24** | **6781.06** | **2106.297** | **- 74.39** | **-0.662** | **80.386** | **<0.001** |
| **ΔBR** | **345.44** | **0.825** | **342.94** | **0.805** | **-2.50** | **-0.020** | **4.502** | **0.034** |
| **ΔSIR** | **386.19** | **0.906** | **380.73** | **0.874** | **-5.46** | **-0.033** | **7.455** | **0.006** |
| **ΔF** | **346.79** | **1.333** | **343.57** | **1.291** | **-3.22** | **-0.042** | **5.224** | **0.022** |
| **ΔITS** | **210.59** | **0.774** | **202.25** | **0.714** | **-8.35** | **-0.060** | **10.348** | **0.001** |
| ΔB | 409.75 | 1.345 | 408.12 | 1.318 | -1.63 | -0.026 | 3.630 | 0.057 |
| Δ16s | 391.20 | 2.724 | 391.46 | 2.718 | 0.26 | -0.006 | 1.742 | 0.187 |
| ΔMBC | 366.06 | 0.771 | 369.51 | 0.771 | 3.45 | -0.001 | 1.447 | 0.229 |
| ΔMBN | 264.01 | 0.519 | 267.69 | 0.517 | 3.68 | -0.002 | 1.679 | 0.195 |
| ΔLC | 157.45 | 0.339 | 162.11 | 0.338 | 4.65 | -0.001 | 2.655 | 0.103 |
| ΔLN | 78.97 | 0.256 | 84.93 | 0.256 | 5.96 | -0.001 | 3.962 | 0.047 |
| ΔRC | 378.78 | 0.670 | 382.58 | 0.669 | 3.80 | -0.001 | 1.806 | 0.179 |
| Δ$qCO_2$ | 365.54 | 367.81 | 367.81 | 1.133 | 2.28 | -0.002 | 0.278 | 0.598 |
| **ΔCUE** | **221.97** | **0.505** | **217.71** | **0.485** | **-4.26** | **-0.020** | **6.265** | **0.012** |
| ΔShoot | 261.17 | 0.334 | 266.09 | 0.333 | 4.92 | -0.001 | 2.918 | 0.088 |
| ΔRoot | 89.52 | 0.248 | 88.454 | 0.243 | -1.07 | -0.005 | 3.065 | 0.080 |

Table S4: Individual pieces of mixed-effects models used in SEM (Fig. 4) to test the hypothesis that grazer-effect on soil-C is mediated by their effect on microbial functions.

Model: ΔF~Assemblage-type, random = ~1|time/sampling locations

| Source | Estimate (mean ± SE) | t-value | P-value |
|---|---|---|---|
| Intercept | 1.00±0.30 | $t_{82}$=3.33 | <0.01 |
| Assemblage-type | **-0.80±0.34** | **$t_{82}$=-2.38** | **0.02** |

Model: ΔMBC~Assemblage-type+ΔF, random = ~1|time/sampling locations

| Source | Estimate (mean ± SE) | t-value | P-value |
|---|---|---|---|
| Intercept | -0.05±0.21 | $t_{73}$=-0.23 | 0.82 |
| **Assemblage-type** | **0.38±0.23** | **$t_{73}$=1.70** | **0.09** |
| ΔF | 0.05±0.07 | $t_{73}$=0.70 | 0.49 |

Model: ΔLC~Assemblage-type+ΔMBC+ΔLN, random = ~1|time/sampling locations

| Source | Estimate (mean ± SE) | t-value | P-value |
|---|---|---|---|
| Intercept | 0.14±0.08 | $t_{63}$=1.69 | 0.10 |
| Assemblage-type | -0.11±0.11 | $t_{63}$=-0.99 | 0.33 |
| ΔMBC | 0.01±0.05 | $t_{63}$=0.11 | 0.91 |
| ΔF | 0.03±0.03 | $t_{63}$=0.96 | 0.34 |

Model: ΔLN~Assemblage-type+ΔMBC, random = ~1|time/sampling locations

| Source | Estimate (mean ± SE) | t-value | P-value |
|---|---|---|---|
| Intercept | 0.18±0.08 | $t_{74}$=2.21 | 0.03 |
| Assemblage-type | -0.04±0.06 | $t_{74}$=-0.69 | 0.49 |
| ΔF | 0.01±0.02 | $t_{74}$=0.47 | 0.64 |

Model: ΔCUE~ΔF+ΔMBC+ΔLC+ΔLN, random = ~1|time/sampling locations

| Source | Estimate (mean ± SE) | t-value | P-value |
|---|---|---|---|
| Intercept | -0.02±0.07 | $t_{45}$=-0.28 | 0.52 |
| **ΔF** | **0.09±0.03** | **$t_{45}$=2.29** | **0.04** |
| ΔMBC | 0.02±0.06 | $t_{45}$=0.27 | 0.79 |
| ΔLC | 0.07±0.41 | $t_{45}$=0.18 | 0.86 |
| ΔLN | 0.01±0.44 | $t_{45}$=0.02 | 0.99 |
